# Automated Quantification of Mitochondrial Fragmentation in an In-Vitro Parkinson’s Disease Model

**DOI:** 10.1101/2020.05.15.093369

**Authors:** Daniel J. Rees, Luke Roberts, M. Carla Carisi, Alwena H. Morgan, M. Rowan Brown, Jeffrey S. Davies

## Abstract

Neuronal mitochondrial fragmentation is a phenotype exhibited in models of neurodegeneration such as Parkinson’s Disease. Delineating the dysfunction in mitochondrial dynamics found in diseased states can aid our understanding of underlying mechanisms for disease progression and possibly identify novel therapeutic approaches. Advances in microscopy and the availability of intuitive open-access software has accelerated the rate of image acquisition and analysis, respectively. These developments allow routine biology researchers to rapidly turn hypotheses into results. In this protocol, we describe the utilisation of cell culture techniques, high-content imaging (HCI), and subsequent open-source image analysis pipeline for the quantification of mitochondrial fragmentation in the context of an in-vitro Parkinson’s Disease model.

## Introduction

The extraction of substantial quantitative data from fluorescence and light microscope images of in-vitro and ex-vivo biological samples is beyond the ability of most day-to-day laboratory researchers. In the last few decades, the development of microscopy technology and the blending of routine biology and computer science skills has resulted in the development of intuitive open source image analysis software to support laboratory experiments. The existence of software allows the successful automation of data extraction processes to liberate investigators from hours of image-by-image manual data capture. As a result, the routine utilisation of such techniques by biology researchers has accelerated (Kamentsky et al., 2011). Many protocols (or pipelines) developed can be readily adapted by biologists to analyse myriad cell biology contexts. Here, we describe a protocol for the analysis of mitochondria.

In eukaryotic cells mitochondria continuously undergo flux between fission and fusion resulting in an observable difference morphology (Scott & Youle, 2010). Dysfunction in mitochondrial dynamics in neurons is recognised as a measurable phenotype of in-vitro and in-vivo models of Parkinson’s Disease (PD) and other neurodegenerative diseases such as Alzheimer’s Disease and Huntington’s Disease (Franco, Li, Rodriguez-Rocha, Burns, & Panayiotidis, 2010; Haque et al., 2008; Pozo Devoto & Falzone, 2017). In-vitro rotenone-based PD models represent a reliable and reproducible tool for investigating disease characteristics through analysis of subcellular organelle morphology (Betarbet et al., 2000; Sherer et al., 2002).

Rotenone is a highly lipophilic compound and an inhibitor of the electron transport system’s complex 1 (NADH Dehydrogenase) (Hoglinger et al., 2005; Jayaraj, Tamilselvam, Manivasagam, & Elangovan, 2013). Inhibition of electron transfer from NADH to ubiquinone in complex 1 of the electron complex system causes a build-up of electrons in mitochondrial inter-membrane space and the increase of the mitochondrial membrane potential (Mehta & Li, 2009). Rotenone exposure induces features of Parkinson’s disease (PD) in-vitro and in vivo, including selective nigrostriatal dopaminergic degeneration and formation of ubiquitin- and α-synuclein-positive inclusions (Betarbet et al., 2000; Sherer et al., 2002).

In our in-vitro PD model, mitochondrial fragmentation is elevated following challenge with the neurotoxin rotenone. The 24hr neurotoxin challenge also results in decreased cell viability, increased cell cytotoxicity and loss of cell number. Here we describe a simple, yet robust protocol for the quantification of mitochondrial fragmentation using routine laboratory neuron cell culture techniques aided by automated analysis via open-source computer software. See figure 12 for supplementary data.

In this paper we separate the protocol into its two key strands: SN4741 Neuron culture and treatment in a Rotenone-based model of Parkinson’s Disease (PD), which outlines an in-vitro rotenone PD model, fluorescence labelling of mitochondria and image acquisition (basic Protocol 1); and a step-by-step construction and execution of an automated CellProfiler™ (CellProfiler) pipeline for the analysis of mitochondrial fragmentation (basic protocol 2).

## Basic Protocol 1: SN4741 Neuron Culture and Treatment in a Rotenone-based model of Parkinson’s Disease

This protocol details wet-lab sample preparation and subsequent image acquisition procedures. The images acquired in this protocol are saved onto a physical or cloud storage platform. A step-by-step guide to build and execute an automated CellProfiler (https://cellprofiler.org/) analysis pipeline for the extraction of mitochondrial measurements will be detailed in *‘Basic Protocol 2’*.

In Basic Protocol 1 we acquire images for subsequent data measurements related to mitochondrial health and fragmentation status. We utilise an in-vitro rotenone-based PD model using which results in increased cytotoxicity, coupled with decreased neuron viability and neuron loss (see Figure 12). At the experimental endpoint, Multitracker Orange (HCS Mitochondrial Health Kit, Invitrogen, Cat. No #H10295) is used to label mitochondria with polarized membranes. In this assay, signal intensity in the orange channel is proportional to membrane potential and mitochondrial health. The experimental set-up was designed for testing the effect of a range of putative anti-PD compounds on mitochondria structure and function.

Membrane potential is a central feature of healthy mitochondria, and membrane depolarization is a good indicator of mitochondrial dysfunction and a measure of cytotoxicity (Jayaraj et al., 2013; Reddy, 2009). This protocol uses Invitrogen MitoTracker Orange as an indicator of mitochondrial function because its accumulation in the mitochondria of live cells is proportional to the mitochondrial membrane potential (Scorrano, Petronilli, Colonna, Di Lisa, & Bernardi, 1999). Invitrogen Hoechst 33342 is used as a segmentation tool to identify cells. Hoechst 33342 nucleic acid stain is a popular cell-permeant nuclear counterstain that emits blue fluorescence when bound to dsDNA. Hoechst 33342 binds preferentially to adenine-thymine (A-T) regions of DNA. This stain binds into the minor groove of DNA (Sandhu, Warters, & Dethlefsen, 1985). A viability stain, such as the Invitrogen Image-iT DEAD Green Viability Stain, can be easily incorporated to further multiplex the assay, however, is omitted from this experiment. These dyes have sufficient retention of fluorescence signal intensity upon formaldehyde fixation and detergent permeabilization to be useful in fixed endpoint assays, as well as applications involving immunocytochemistry for specific protein detection.

Briefly, SN4741 neurons, an immortalised midbrain-derived dopaminergic mouse cell line (Son et al., 1999), were treated 24hs with rotenone, then incubated with MitoTracker Orange, as described in manufacturer ‘s user manual, prior to formaldehyde fixation and Hoechst 33342 nuclear counterstaining. We subsequently use a CellProfiler image analysis pipeline to quantify intracellular mitochondrial fragmentation and fluorescence intensity of mitochondria in the peri-nuclear area of SN4741 neurons-details of subsequent step are in basic protocol 2.

*Note: Herein, MitoTracker Orange will be referred to as ‘MitoHealth’.*

### Materials

Toshiba HDTB320EK3AA 2 TB Canvio Basics USB 3.0 Portable External Hard Drive (Toshiba)

Microscope system: HCA wide-field fluorescence (GE Healthcare InCell 2000/6000), x40 Objective lens. To acquire 52 fields of view per well.

MitoTracker Orange and Hoechst 33342 from the Invitrogen HCS Mitochondrial Health Kit (Invitrogen, #H10295)

SN4741 mouse midbrain derived neuronal cell line cryopreserved in liquid N2.

SN4741 cell culture media

DMEM - Dulbecco’s Modified Eagle Medium (high-glucose (4.5g/L) DMEM without sodium pyruvate (Invitrogen, 41965-039) supplemented with10% filtered FBS, 1% Penicillin/Streptomycin, 1% (100x) L-glutamate and 3% glucose (20% solution)
50mL FBS (10106169, Invitrogen)
5mL Penicillin/Streptomycin (Pen/Strep) (15140122, Gibco, UK)
5mL L-Glutamine (25030081, Gibco, UK)
15mL 20% Glucose solution, filtered bioultra for molecular biology 20% in water (Sigma, 49163-100mL)

T75 cell culture flasks (156499, Thermo Fisher)

Corning® 96-well Clear Flat Bottom Polystyrene TC-treated Microplates, Individually Wrapped, with Lid, Sterile (3596, Corning)

Benchtop vortex

100mM rotenone (Tocris, Cat. No. 3616) stock diluted in Dimethyl Sulfoxide (DMSO) (Sigma-Aldrich, D2650). 50 mg in 1.267mL DMSO.

Trypsin/EDTA (15400054, Gibco, UK)

Paraformaldehyde, 16% w/v aq. soln. (Fischer Scientific)

Water bath set to 37.5 °C for pre-warming cell culture media prior to application

P1000, P200, P20 Pipettes and filtered-pipette tips (Fisherbrand™)

Cell culture incubator. Set at 37°C and 5% CO_2_.

25mL, 10mL and 5mL Sterile Pipettes. Individually wrapped (Fisherbrand™)

Phosphate Buffered Saline PBS (pH 7.4) (10010023, Gibco, UK)

Countess™ Automated Cell Counter (Cat no. C10227, Invitrogen) or manual haemocytometer

Trypan blue stain (0.4%) (15250061, ThermoFisher)

### Seeding SN4741 cells for In-Vitro Rotenone PD Model

1. Retrieve cryovial of SN4741 cells (approximately 1×10^6^ cells) from Liquid nitrogen storage liquid.
2. Thaw a cryovial of SN4741 cells in water bath and pipette contents into 20mL of pre-warmed full cell culture medium to expand culture for experimental procedure. Culture SN4741 cells in complete medium in T75 cell culture vessel under standard and sterile conditions until flask is 80-90% confluent. SN4741 cells require approximately 16hrs to settle and adhere. Cell culture medium is replaced the day after revival (16hrs) following 1×10mL PBS wash. Cell medium should be replenished every 2-3 days. SN4741 neurons in T75 flasks should reach 80-90% confluency in 3-4 days.
3. Remove excesses medium from T75 flask and wash once with 10mL PBS (without Calcium and Magnesium). Swirl flask and use pipette to remove excess.
4. Apply 3-4mL of Trypsin/EDTA solution to culture vessel; swirl to ensure coverage and return flask to incubator for 3-5mins until SN4741 cells become detached from surface of culture vessel. Check flask every few minutes to gauge cell detachment. Gently tap flask to detach residual cells from surface of the T75 flask.
5. Neutralise the cells in suspension with the addition of 4mL cell culture medium.
6. Using a sterile stripette, pipette cell suspension up and down against surface of flask (∼10 times) to ensure single cell suspension and removal of the remaining adhered cells.
7. Transfer cell suspension to 15mL tube and perform cell count and viability test using trypan blue and countess (automated) or haemocytometer (manual) scoring.
8. Prepare a cell suspension with a cell concentration of 30,000 cells/mL (i.e. 3,000 cells/100uL).
9. Using a P200, pipette 100µL (∼3,000 cells) of the cell suspension into each experimental well of a 96-well plate.
10. Place plate in incubator overnight (16-24hrs) for cells to adhere and proliferate.
11. Add 100µL of culture media into each well following 24hrs incubation and incubate for a further 24hrs.

### Preparation and Addition of 10nM Rotenone in In-Vitro PD model

Rotenone stock is prepared by the addition of DMSO to a concentration of 100mM. Aliquots are prepared and sealed into a 50mL falcon tube, protected from light and stored at −20°C.

12. Dilute 50mg of powdered rotenone in 1.267mL DMSO to prepare 100mM rotenone stock. Stock is aliquoted into microcentrifuge tubes, protected from light and stored at 20°C.
13. Thoroughly mix (vortex) 3µL of 100mM rotenone stock diluted in DMSO into 10mL culture medium producing a 30,000nM rotenone solution.
14. Sequentially prepare 3,000nM, 300nM and 30nM rotenone solutions by performing serial 1 in 10 dilutions i.e. 100µL of 30,000nM rotenone solution mixed into 900 µL of cell medium to produce 1mL of 3,000nM rotenone solution etc.
15. Vehicle treatment was prepared from DMSO using the same dilution method.
16. Add 100µL 3x concentrated rotenone solution (30nM) to appropriate wells (this is a 1:3 dilution resulting in final well concentration of 10nM).
17. Incubate cells for 24hr under standard conditions (5% CO_2_; 37°C).
  Note: 100µL of a 50:50 solution of DMSO in PBS was used as a positive control for analysis of cell death.

#### Application of Mitohealth Dye and Image Acquisition at Experimental Endpoint

MitoHealth solution was prepared according to the kit developers’ instructional protocol. Mitohealth kit contains Hoechst (Blue) and Mitohealth stain (red/orange).

18) Remove 175µL of culture media from appropriate wells. Leaving a reminder of 125µL.
19) Add 1.75µL Mitohealth solution into 1mL of medium. *Protect from light.*
20) Add 50µL of staining solution prepared in step (2) to each well. *There should now be 175*µL *of solution in each well.*
21) Incubate cells under normal cell culture conditions for 30 mins.
22) Prepare fixation-counterstaining solution by adding stock reagents 16% PFA and Hoechst 33342 to PBS (without MgCl2 and CaCl2) in the ratio 3mL: 6µL: 9mL. This makes a total of 14mL of fixation-staining solution containing 4% PFA and 1:2000 Hoechst 33342.
  Example: 300µL 16% PFA: 0.6µL Hoechst 33324: 900µL PBS; yields 1200µL of fixation-staining medium.
23) Remove cell culture medium.
24) Add 100µL of fixation-staining solution to each well. Protect from light and incubate at room temperature for 15mins.
25) Remove fixation-staining solution and gently wash cells once with 100µL PBS. Thus, removing excess staining solution.
26) Add 200µL PBS into each well.
27) Proceed to **image acquisition.** Acquire 52 x40 magnification 2D-Deconvoluted .tif format images per well. *Table 1 summarised excitation and emission filters required for image acquisition.*
28) Store image files on external hard drive for downstream image analysis with CellProfiler image analysis pipeline (*described in Basic Protocol 2*).

**Table 1.** Summary of dyes and filters required for image acquisition.

| **HCS Mitochondrial Health Kit stain** | **Approximate FI Excitation Wavelength** | **Approximate FI Emission Wavelength** | **Location of cellular stain** | **InCell x40 objective filter** |
| --- | --- | --- | --- | --- |
| MitoHealth stain | 550nm | 580nm | Perinuclear/<br>Cytoplasmic | Cy3 |
| Image-iT® DEAD Green™ Viability Stain | 488nm | 515nm | Nucleus | FITC |
| Hoechst 33342 | 350nm | 461nm | Nucleus | DAPI |

## Basic Protocol 2: Identification of Cell Nuclei, measurement of Mitochondrial Membrane Potential and Mitochondrial Fragmentation in Mouse Derived Midbrain Dopaminergic Neurons

CellProfiler™ software is a flexible, user-friendly platform on which to conduct image analysis. The software and corresponding website -http://www.cellprofiler.org <u>(RRID:SCR_007358)</u>-contain pre-made computer algorithms (modules) and example analysis pipelines, respectively, CellProfiler is not an out-of-the-box, plug-and-play image analysis software. Rather, these pre-made modules and example pipelines are used to teach researchers, with little computer programming or macro script writing experience, how to design and assemble an image analysis pipeline to fit the specific needs of their experimental question. Interestingly, CellProfiler also allows for ImageJ (NIH) software input of macro scripts to aid image analysis.

The protocol below describes the step-by-step CellProfiler pipeline for the identification of nuclei, measurement of MitoHealth staining intensity and quantification of mitochondrial fragmentation protocol used to quantify SN4741 cell mitochondrial measurements from x40 magnification digital images. This protocol was developed by optimising a range of modules and example protocols. Each step includes a description, CellProfiler-derived notes and rationale followed by instructions on how to execute the step in CellProfiler software (*in italic font).* ‘Notes’ are referenced from CellProfiler.org/manuals. The CellProfiler pipeline and modules described here is transferable between CellProfiler software versions 2.0 and 3.0 (Kamentsky et al., 2011; McQuin et al., 2018).

### Materials

Digital images: on internal/external hard drive or cloud storage.

Laptop or desktop PC: macOS, Windows (32-bit) or Windows (64-bit).

MS Excel software

GraphPad Prism for graphs and statistical analysis

Toshiba HDTB320EK3AA 2 TB Canvio Basics USB 3.0 Portable External Hard Drive for data storage.

Cell Profiler software: version 2.1.0 or newer. Cell profiler versions 2.2.0 or newer will require Java 8. Available at: https://cellprofiler.org/releases/

### Step 1: Upload Image Set, Image Sorting and Image Pre-Processing and Illumination Correction

1. Open Cell Profiler Software and select the experimental image set.
2. *Open Cell Profiler > Select ‘Images’ in ‘Input Module’ window > Right click inside empty ‘File list’ window > Browse and select image set.* Here, our image set is composed of two channels, Cy3 and DAPI, which account for x40 magnification images of MitoHealth stained mitochondria and Hoechst stained nuclei of SN4741 cells respectively (see figure 2 for illustration). To filter the images according to filename:
3. *Select ‘Names and Types’ in pipeline Input modules > Select ‘Images Matching Rules’ then select ‘File’, ‘Does’, ‘Contain’ from dropdown box selections > Type ‘DAPI’ into input box to group DAPI channel images.*
4. *Select ‘Add another image’ and repeat the process with exception to entering ‘Cy3’ into input box to group Cy3 channel images. Assign a (user-defined) name for grouped files > Select ‘Update’.* **Note:** Care must be taken when filtering image sets by common phrases in filenames, such as ‘DAPI’ and ‘Cy3’, as phrases are case-sensitive. Inconsistency in spelling or case-sensitivity results in failure to identify files and sub-group images. Experimental image set should now be grouped in two file lists under user-defined group titles e.g. DAPI and Cy3 and ranked in numerical order.

During the analysis of images taken with a fluorescence microscope it is common for the images to be pre-processed prior to analysis. The most common reason for this is uneven illumination across a field of view as the light source does not evenly illuminate across the entire area. This results in areas with greater illumination than other areas, usually the corners of the image.

Calculate correct illumination function:

5. *Select* ***Correct Illumination Calculate*** *module > Select input image (name assigned from Names and Types).*
6. *Insert a (user-defined) Name the Output Image e.g. ‘IllumCy3’.*
7. *Select ‘Regular’ option in How the illumination function is calculated.* **Note**: The option ‘Regular’ is selected if stained objects (e.g. cell bodies, nuclei) are evenly dispersed across your image(s) and cover most of the image. This is true in the case of Cy3 and DAPI channel images. Regular intensities make the illumination function based on the intensity at each pixel of the image (or group of images if you are in ‘All’ mode) and applied by division using ***Correct Illumination Apply* module.**
8. *Select ‘Median Filter’ > Select ‘Manual’ method to calculate filter size > input smoothing filter size ‘325’.* **Note:** Illumination function was calculated across the entire image set prior to initiation of object identification and subsequent data measurements.

To apply the illumination function calculated by ***Correct Illumination Calculate***:

9. *Select* ***Correct Illumination Apply*** *module.*
10. *Input > a user defined name into ‘Name the output image’ box e.g. ‘CorrCy3’.*
11. *Select illumination function from drop down box (as defined in* ***Correct Illumination Calculate*** module *e.g. IllumCy3).*
12. *Select ‘Divide’ as method for how the illumination function is applied.* **Note:** Divide is the recommended method of illumination function application by CellProfiler users if the illumination correction function was created using ‘Regular’ in the ***Correct Illumination Calculate*** module.

Figure 2 illustrates the shading seen on edges of Cy3 and DAPI channel images acquired images before and after application of illumination function as described above.

**Figure 1:**
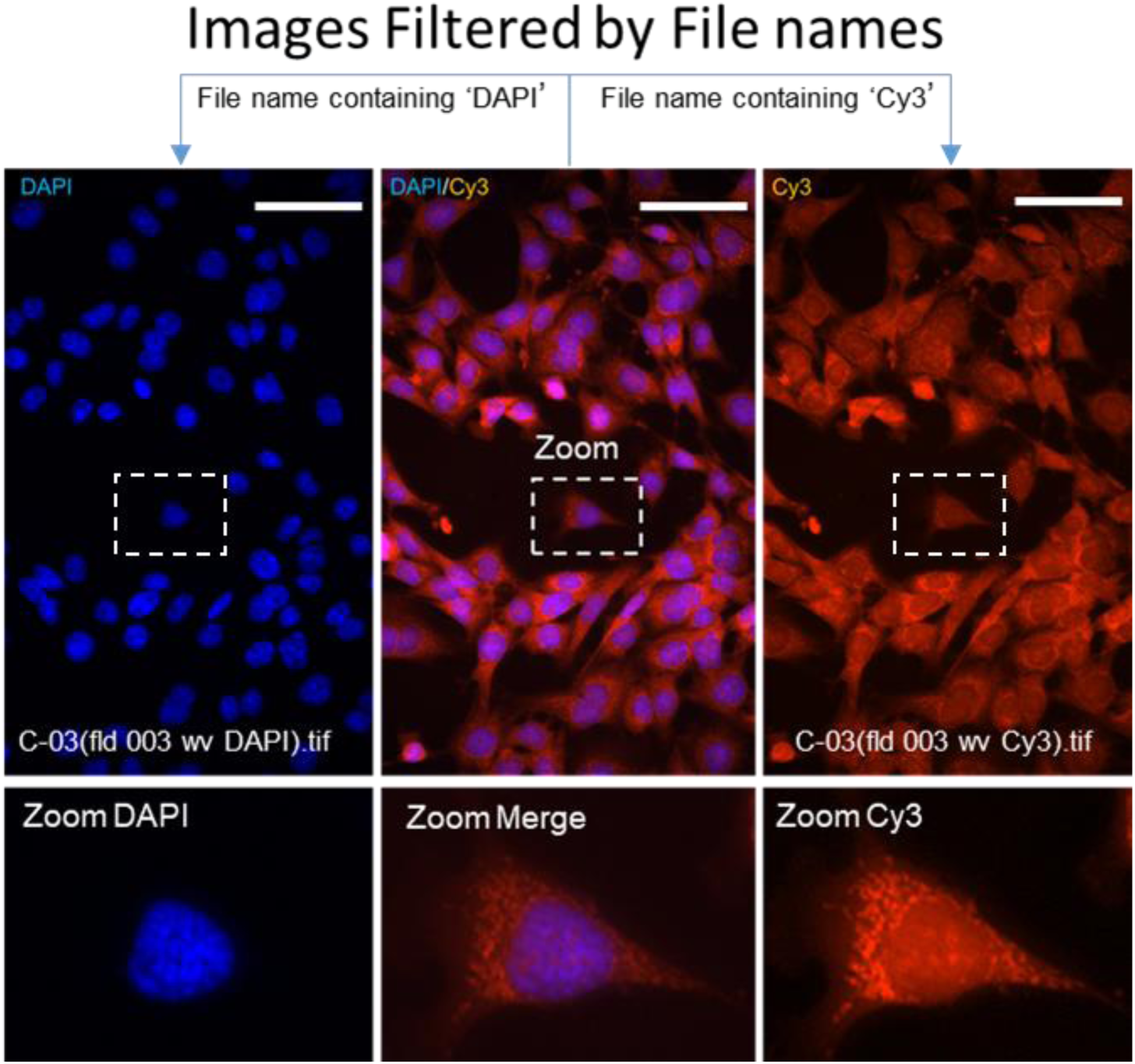
Images are Filtered into Groups by File Names. Figure 1 illustrates an image set containing DAPI and Cy3 images being separated into groups (DAPI and Cy3) based on their file names. Cropped image tile labelled DAPI/Cy3 represents a pseudo-coloured composite of both channels. Blue arrows illustrate the separation of DAPI and Cy3 channels based on contents of their file names. Example image filenames are given on pseudo-coloured DAPI and Cy3 images. Each pseudo-coloured image has a representative enlarged image for graphical purposes (dotted white box). Scale bar = 40µm.

**Figure 2:**
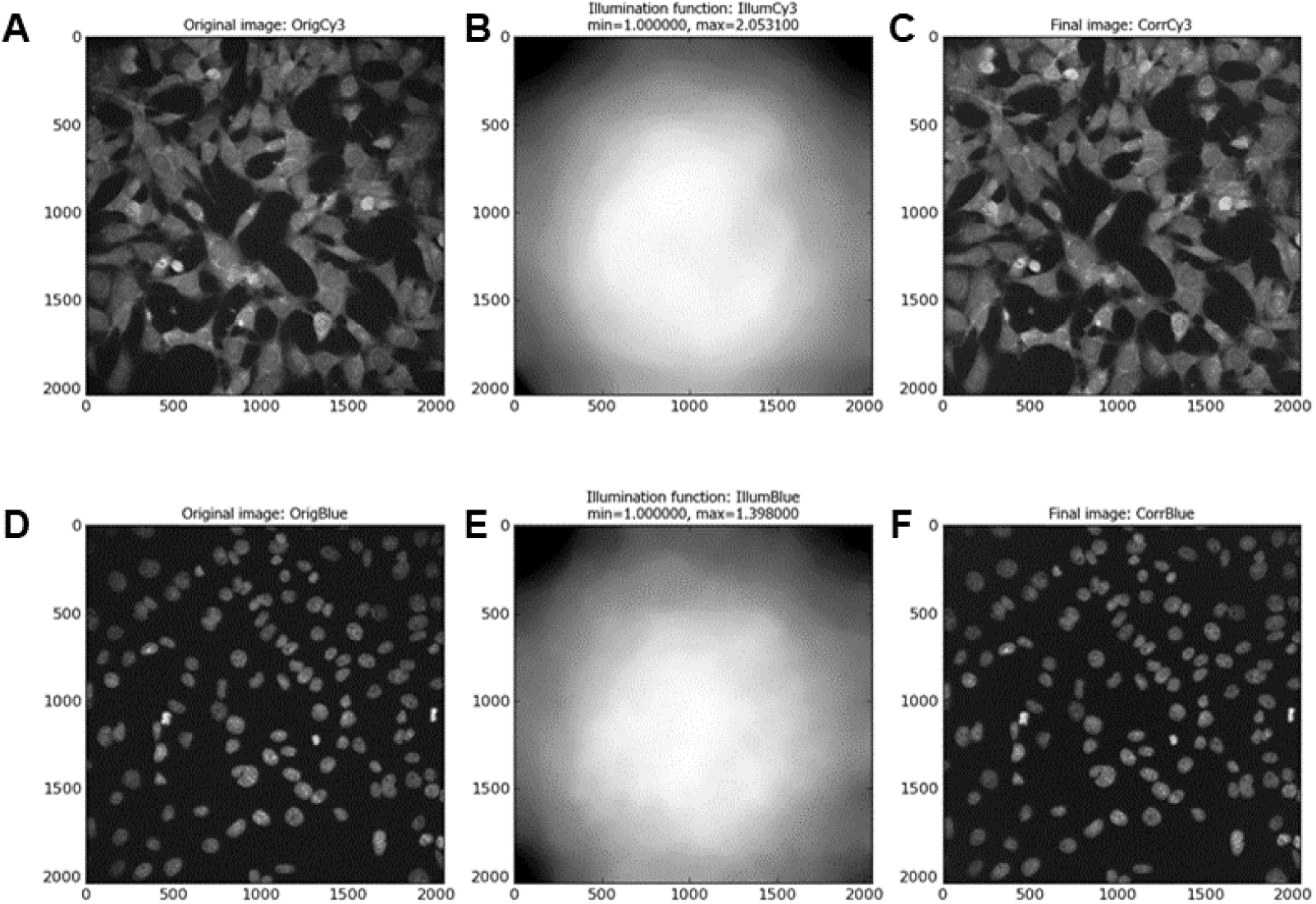
Cy3 Image and DAPI Channel Images Before and After Illumination Correction. Figure 2 (**A**) and (**C**) show representative x40 magnification greyscale images acquired using In Cell 2000 Cy3 filters before (OrigCy3) and after (CorrCy3) illumination correction, respectively. Figure 2 (**B**) illustrates the illumination function (IllumCy3) produced from entire Cy3 image set. Figure 3 **D** & **F** show representative x40 magnification greyscale images acquired using In Cell 2000 DAPI filters before (OrigBlue) and after (CorrBlue) illumination correction, respectively. Figure 2 (**E**) (IllumBlue) illustrates the illumination function produced from entire image set. This illumination correction is carried out by CellProfiler modules CorrectIlluminationCalculate and CorrectIlluminationApply. Images (**A**-**F**) are micro pictograms acquired and saved from the CellProfiler Software.

### Step 2: Identification of Nuclei

Here we identify cell nuclei (a feature) based on their high contrast background to foreground staining intensity. In CellProfiler software, a primary object is defined as an object or feature (usually a cellular sub-compartment e.g. nucleus) which can be used as the basis on which the subsequent image analysis steps can be built upon.

Nuclei are commonly selected as the primary object of an image analysis pipeline due to their relatively uniform morphology and high background to foreground staining intensity ratio which makes them easily identifiable.

13. *Select >* ***‘Identify Primary Objects’*** *module.*
14. *Input > (user named illumination corrected) DAPI image(s).*
15. *Input > Typical Nucleus diameter range (pixels). Min: 50 and Max: 135.*
16. *Discard objects outside the diameter range > Yes.*
17. *Discard objects touching the border of the image > Yes.* **Note:** Estimate nucleus diameter using CellProfiler by selecting a DAPI image by ‘Right click’ on DAPI image and selecting ‘Show Selected Image’; and selecting ‘Tools > Measure length’.
18. *For thresholding select ‘Global’ Thresholding Strategy with ‘Otsu’ method and ‘Two Classes’ thresholding. Set threshold correction factor of ‘1.0’; and lower and upper bounds to ‘0.0’ and ‘1.0’, respectively.* **Note:** Global thresholding strategy uses the pixel intensities in each un-masked image to calculate a single threshold value to classify pixels as foreground (intensity above the threshold value) or background (intensity below the threshold value). This thresholding strategy is fast and robust and commonly applied to images that have a uniformly illuminated background. Foreground-to-background pixels are easily distinguishable using Hoechst counterstained SN4741 cell nuclei. Foreground and background pixels of Hoechst counterstained nuclei are easily identifiable in DAPI channel images, this we use ‘Global’ thresholding strategy for primary object (nuclei) identification. Otsu is a method of automatically finding thresholds by splitting image pixels into two (foreground and background) or three classes (foreground, mid-level and background) by minimizing the variance within each class. Lower bounds on threshold can be set at 0.0 as there is an object (Nucleus) in each image. Otherwise, an empirically determined lower threshold bound as automatic methods of thresholding may miss-assign pixels in a blank image to the foreground class, potentially yielding false-positive results.
19. *Select > ‘Shape’ for de-clumping of primary object which are touching or in close proximity and ‘Shape’ for drawing dividing lines between objects.*
20. *Select > ‘Yes’ to ‘Automatically calculate size of smoothing filter for de-clumping’.*
21. *Select > ‘Yes’ to ‘Automatically calculate minimum allowed distance between local maxima’.*
22. *Select > ‘No’ to ‘Retain outlines of the identified objects.*
23. *Fill in identified objects > ‘After both thresholding and de-clumping’.* **Note:** By selecting ‘shape’ in CellProfiler pipeline as the de-clumping method, a line will be drawn between the areas of two the nuclei whose boundaries intercept. An intersecting line will be drawn between the two (or more) clumped nuclei and these will then be identified as distinct nuclei. Examples of this are highlighted by white arrows in Figure 3D.

### Step 3: Identify Secondary Objects: Ring shaped Cell Body

The next module in CellProfiler pipeline is ***Identify Secondary Objects*.** This module identifies objects by using the object identified of another module as a starting point. In this example, the primary object (nuclei) identified in the Identify Primary Object will act as the basis for the identification of SN4741 peri-nuclear area (doughnut shaped ring around the nucleus) in the corresponding Cy3 images. This will be executed by dilation of the previously identified nuclei - this dilated nucleus area is named ‘Full Doughnut’ in Figure 4. A subsequent module in this pipeline will remove the nucleus, and a ring like object will be remaining (a doughnut shaped-perinuclear region). This is referred to as ‘True Doughnut’ in Figure 5. Other modules will collect measurements such as area and intensity of peri-nuclear regions.

**Figure 3:**
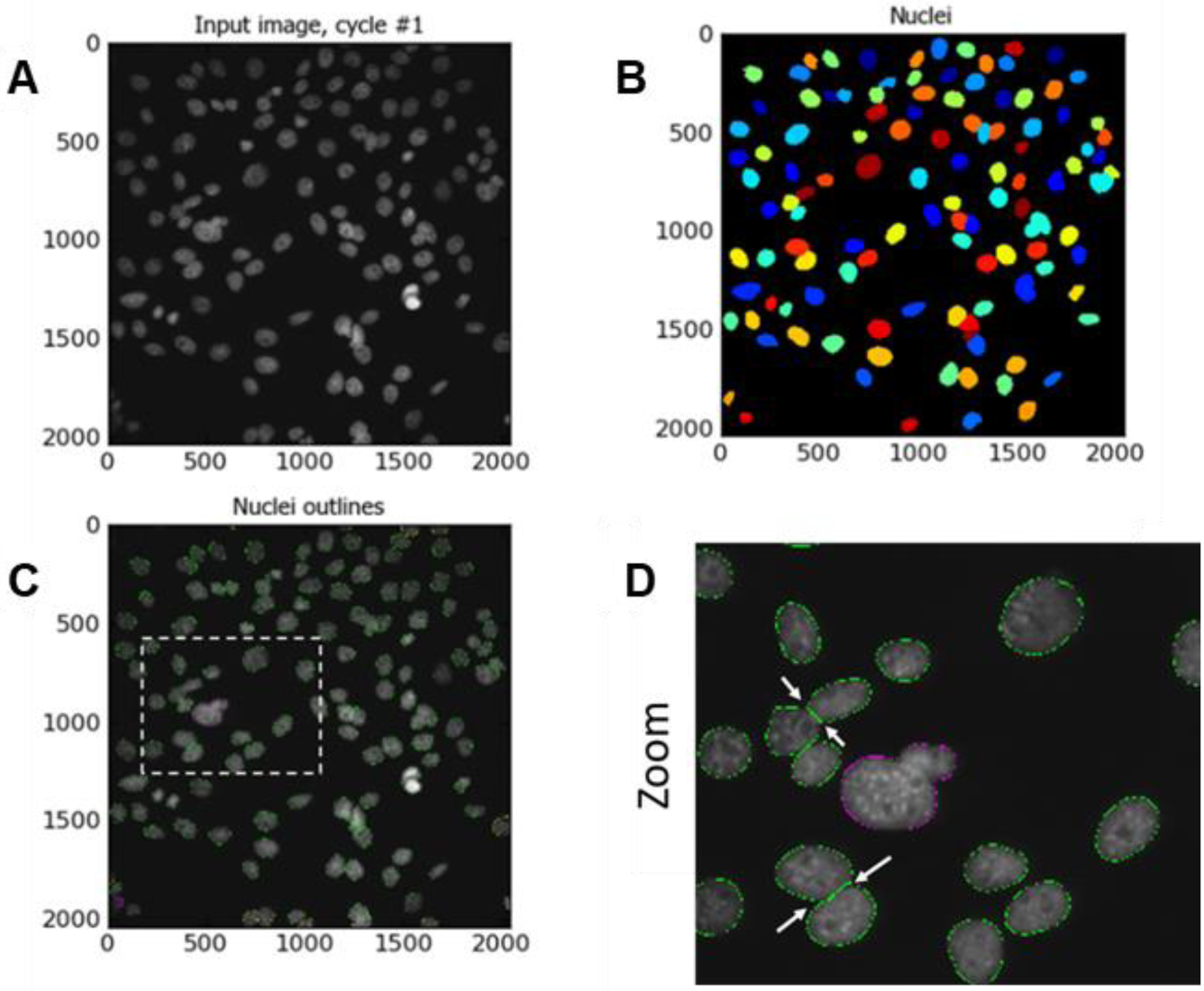
Identify Nuclei as Primary Objects. Figure 3 (**A**) shows a representative illumination corrected x40 magnification image of Hoechst counter-stained nuclei acquired using In Cell Analyser 2000 DAPI channel filters. (**B**) illustrates identified primary objects using object coloured masks. (**C**) shows nuclei fitting to protocol input criteria (outlined in green); nuclei filtered out by size criteria (outlined in magenta); and nuclei filtered out for touching the image border (outlined in yellow). Dashed white box shows enlarged area (**D**). (**D**) shows the enlarged area with white arrows indicate indentations used to de-clump objects (nuclei). Images (**A**-**D**) are micro pictograms acquired and saved from CellProfiler.

**Figure 4:**
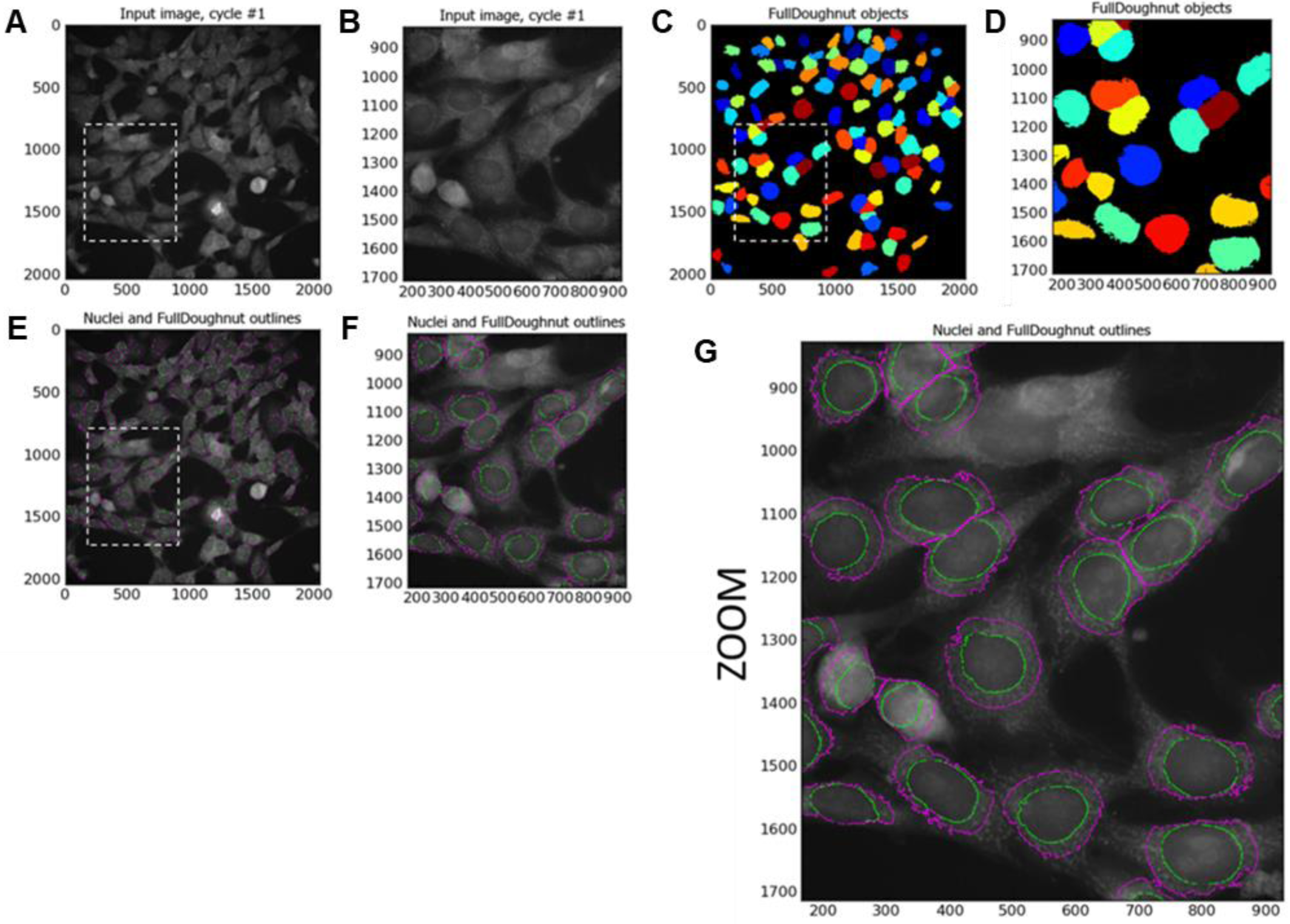
Identification of Secondary Object-Full Doughnut. Figure 4 shows the Illumination Corrected Cy3 image (**A**); the result of a 25-pixel distance dilation in nucleus size with respect to Cy3 image staining intensity thresholding. Identified secondary objects/Full Doughnuts are illustrated in the form of coloured masked objects. Dashed white line boxes in (**A**), (**C**) and (**E**) highlight the enlarged areas illustrated in (**B**), (**D**) and (**F**), respectively. These zoomed images show the image segmentation more clearly. Image (**G**) is a larger version on (**F**), illustrating the nuclear area/primary object with a green outline and the edges of the Full Doughnut/secondary object with a magenta outline. Images (**A**-**G**) are micro pictograms acquired and saved from the CellProfiler Software.

**Figure 5:**
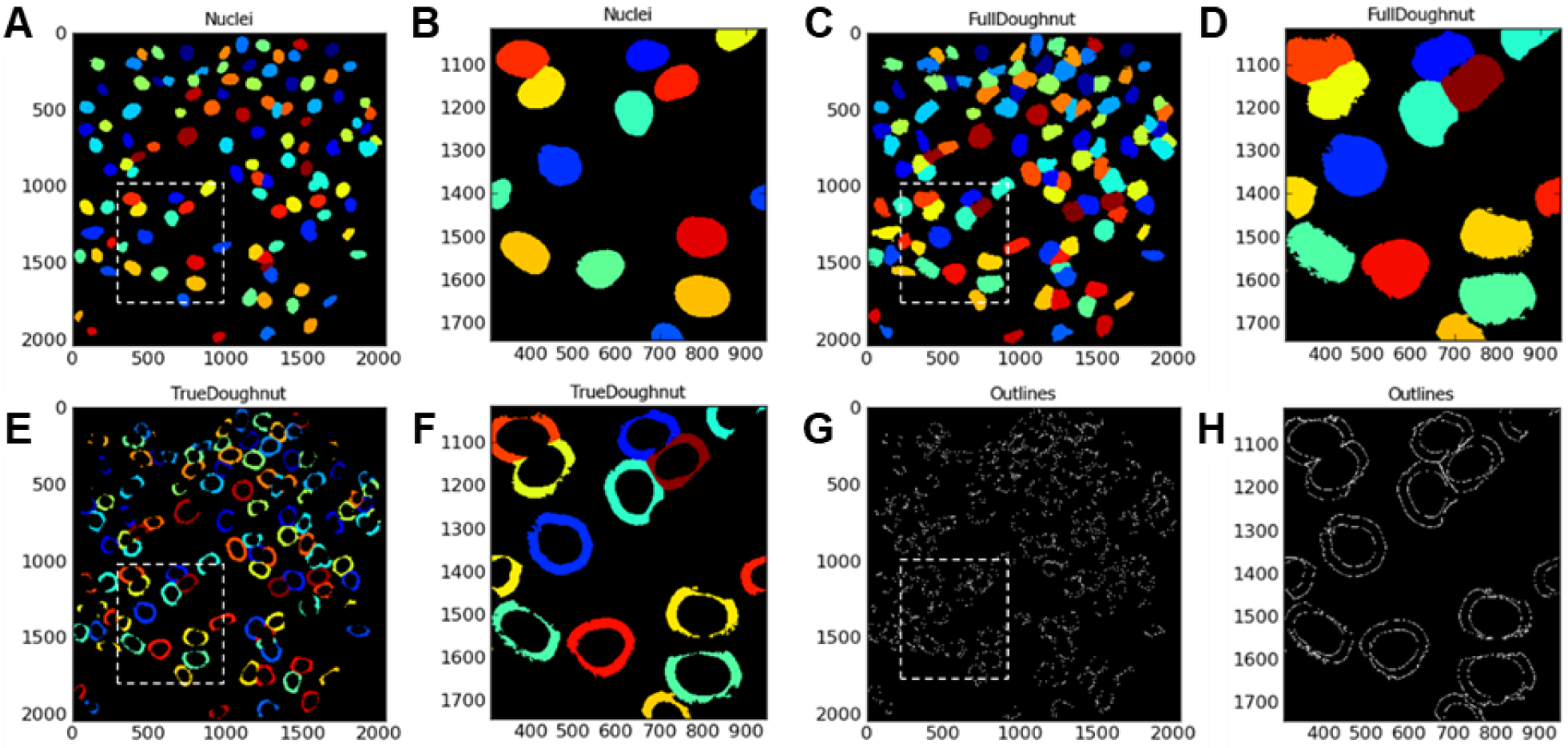
Identify Tertiary Objects. Images (**A**), (**C**) and (**F**) illustrate the masked image result of sequential steps/modules from identification of primary object (Nuclei), identification of secondary object (Full Doughnut) and tertiary object (True Doughnut), respectively. The ring-shaped True Doughnut is the result of removing the Nuclei areas from the Full Doughnuts. Micro-pictograph (**G**) highlights the outlines of the resulting True Doughnut. Dashed white box in images (**A**), (**C**), (**E**) and (**G**) indicated the areas which have been enlarged for illustrative purposes and are represented by images (**B**), (**D**), (**F**) and (**H**), respectively. Images (**A**-**H**) are micro pictograms acquired and saved from the CellProfiler software.

**Figure 6:**
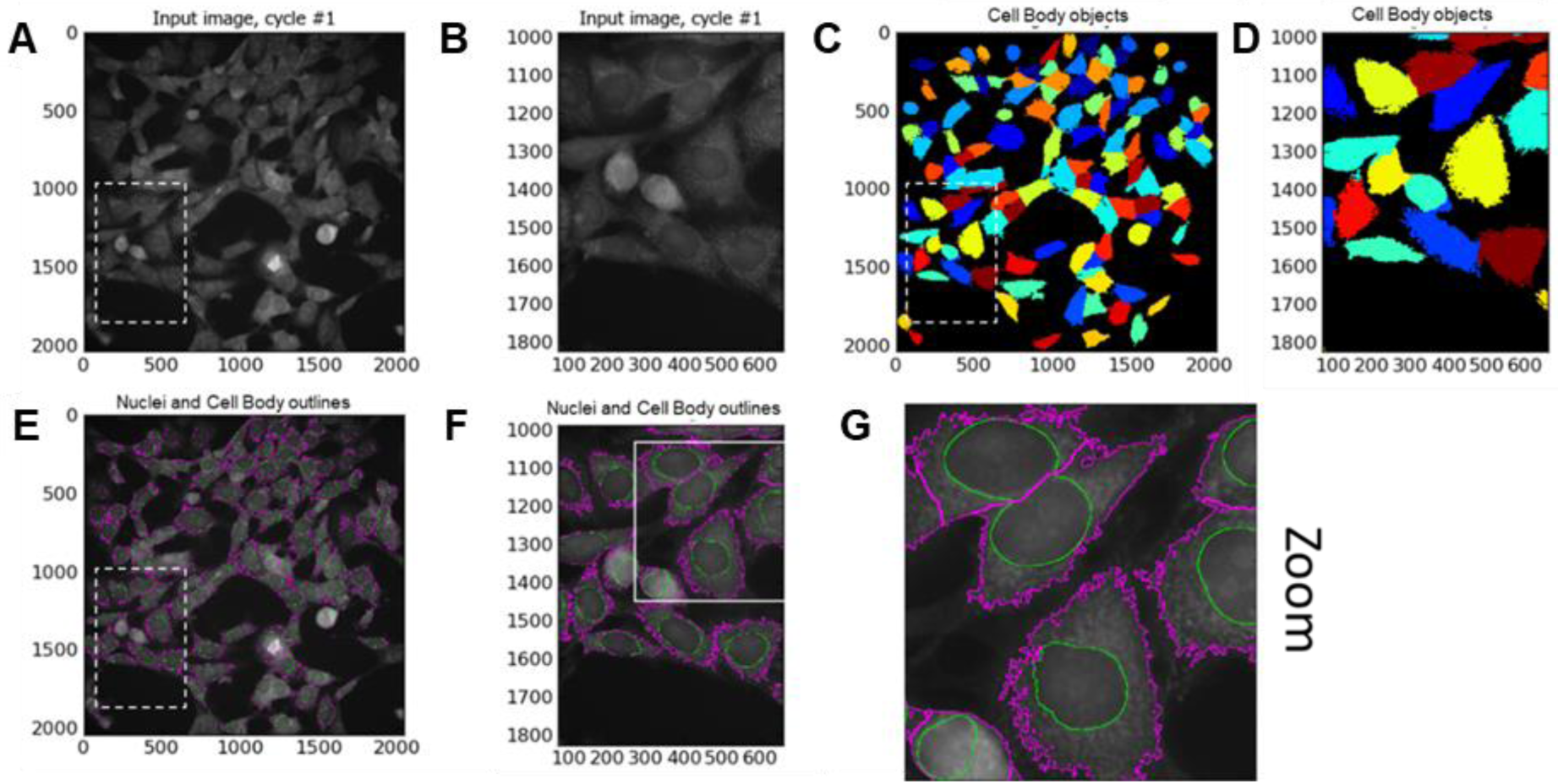
Identify SN4741 Cell Body as Secondary Object. Figure 6 (**A**) shows the input Illumination Corrected x40 magnification Cy3 channel image. (**C**) Illustrates the result of propagation method of identifying cell (body) edge with respect to Cy3 image staining intensity thresholding. Identified secondary objects/Cell bodies are illustrated in the form of coloured masked objects. Image (**E**) shows the input Cy3 channel image with identified cell body outlines in magenta and nuclei outlines in green. Dashed white line box in (**E**) highlights the area enlarged to produce image (**F**). White box in image (**F**) indicates enlarged area to produce image (**G**). Images (**F**) and (**G**) were generated to illustrate image segmentation. Dashed white line boxes in (**A**) and (**C**) highlight areas that are enlarged and illustrated in (**B**) and (**D**), respectively. Images (**A**-**G**) are micro pictograms acquired and saved from the CellProfiler software.

**Figure 7:**
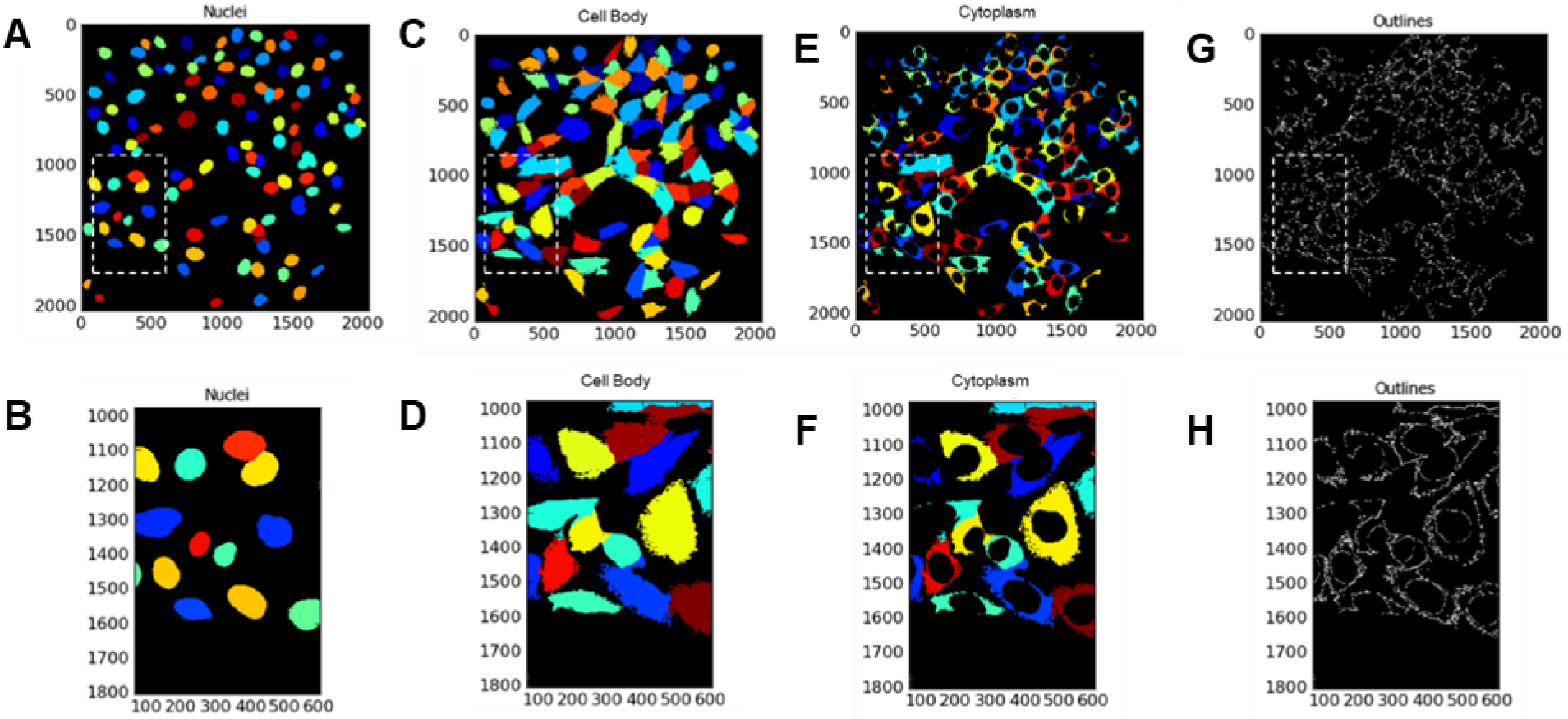
Identify SN4741 Cytoplasm as Tertiary Object. Images (**A**), (**C**) and (**F**) illustrate the masked image result of sequential steps/modules from identification of primary object a (Nuclei), identification of secondary object (cell body) and tertiary object f (cell cytoplasm), respectively. SN4741 cytoplasm is the result of removing the Nuclei areas from the areas of entire cell bodies. Micro-pictograph (**G**) highlights the outlines of the resulting SN4741 cell cytoplasm. Dashed white boxed in images (**A**), (**C**), (**E**) and (**G**) indicated the areas which have been enlarged for illustrative purposes and are represented by images (**B**), (**D**), (**F**) and (**H**), respectively. Images (**A**-**H**) are micro-pictograms acquired and saved from the CellProfiler software.

**Figure 8:**
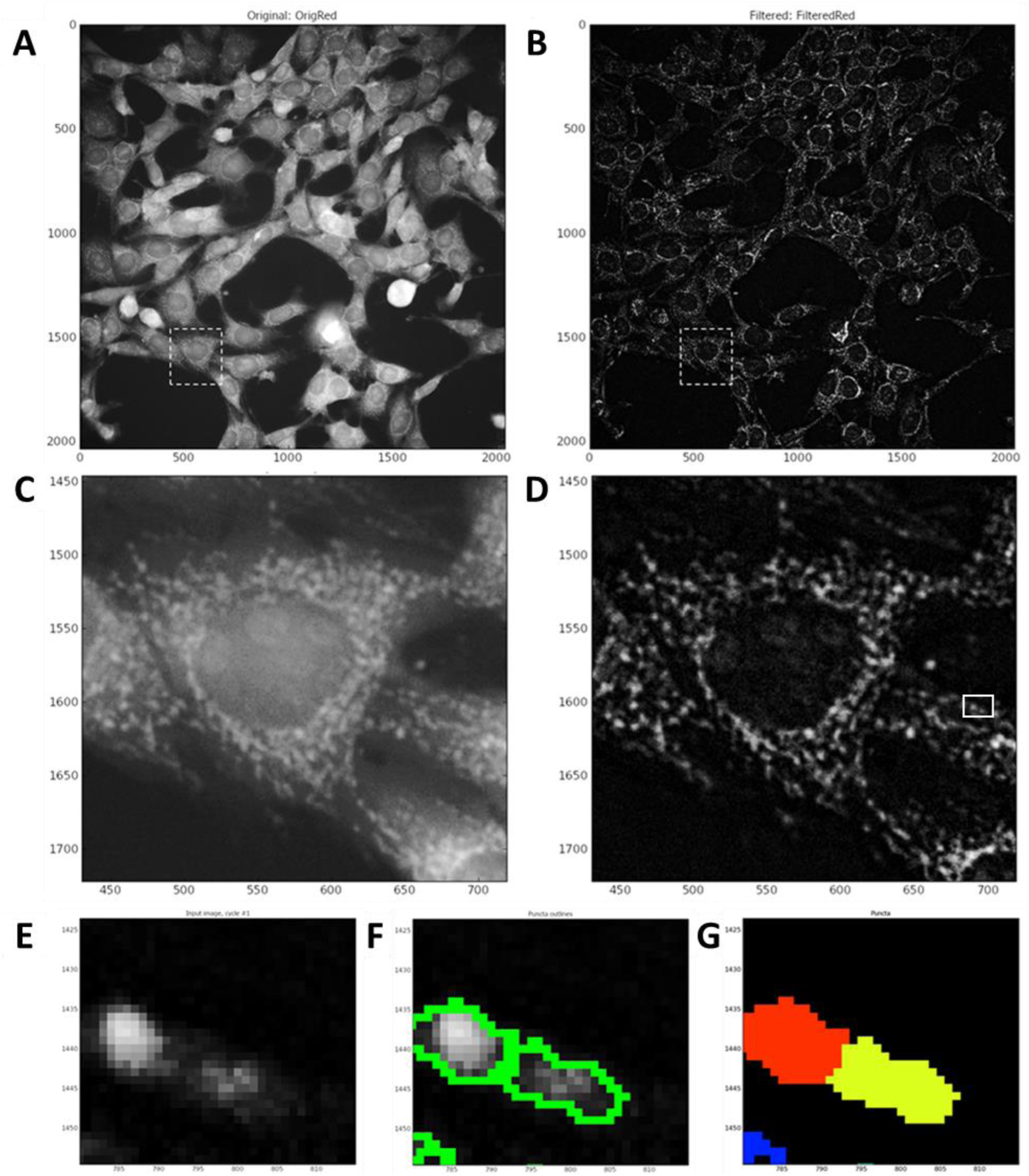
Enhance Mitochondrial Fragments and Identification. Figure 8 (**A**) shows an input ‘CorrCy3’ image prior to speckle enhancement. (**B**) shows image (**A**) post-enhance module. Dashed white line box in (**A**) and (**B**) highlight the area displayed in (**C**) and (**D**) which are pre- and post-enhancement module, respectively. (**E**) illustrates enlarged input image outlined in solid white box in (**D**); these illustrate outlined mitochondrial puncta (**F**); and marks of identified objects (**G**) seen in enhanced image (**E**).

To expand nucleus and create ‘Full Doughnut’ shape:

24. *Select > ‘****Identify Secondary Object’*** *module.*
25. *Input > Illumination function corrected Cy3 image ‘‘CorrCy3’.*
26. *Input (primary) objects > Select ‘Nuclei’. These were identified by Identify Primary Object module.*
27. *Insert the name of objects to be identified > ‘FullDoughnut’.*
28. *Select > ‘Distance-B’ as method to identify the secondary objects.* **Note:** By selecting Distance as the method to identify the secondary objects (‘Full Doughnut’) the primary object (nucleus) is expanded to identify a ‘doughnut’ (or ‘annulus’) shaped region in the cell, usually in the cytoplasm. CellProfiler provides two methods of identifying the secondary object. The first, ‘Distance-N’ does not utilise staining intensity in the image of the second stain - in this instance Cy3. This entails expanding the nucleus a set distance. However, this may include areas of image background as well as cell cytoplasm. Our protocol on the other hand utilises ‘Distance-B’ method. This second ‘Distance’ method expands the nucleus a set distance however, this method uses the thresholding of the secondary staining image to eliminate background regions. Thus, the nucleus is only expanded into areas of foreground staining intensity to identify ‘Full Doughnut’ secondary object without including background image regions.
29. *Select > ‘Automatic’ threshold method.*
30. *Input > ‘25’ as number of pixels to expand the primary object (nucleus).*
31. *Fill holes > Select ‘Yes’.*
32. *Discard Objects touching the border of the image > Select ‘Yes’.*
33. *Retain outlines of the identified secondary objects > Select ‘No’.* **Note:** the ‘Automatic’ thresholding method is the default setting in CellProfiler and is robust. As this strategy is automatic it does not allow users to select the threshold algorithm or to apply additional corrections to the threshold. The ‘Automatic’ strategy calculated the threshold using maximum correlation thresholding (MCT) for the whole image. The threshold is then applied to the image and smoothed with a Gaussian filter with a sigma of 1. This blurs or obscures smaller than an entered diameter and spreads bright or dim features larger than the entered diameter. The MCT method is described by Padmanabhan, Eddy, & Crowley (2010) as computationally efficient and accurate without relying on assumptions of the statistics of the image. This algorithm has been trialled and tested on neuroscience images in the presence and absence of illumination correction to accurately aid in automated image analysis (Padmanabhan et al., 2010).

### Step 4: Identify Tertiary Objects: Cell Cytoplasm or True Doughnut

Following the identification of the ‘Full Doughnuts’ (secondary objects), the ***Identify Tertiary Objects*** module is then used to remove the smaller objects (nuclei) from the larger secondary objects (Full Doughnuts) to leave a ring shape around the nucleus, referred to here as ‘True Doughnut’.

To identify tertiary objects (ring shaped ‘True Doughnuts’):

34. *Select >* ***‘Identify Tertiary Objects’*** *module.*
35. *Select > ‘FullDoughnut’ as larger identified objects (Output from Identify Secondary Object module).*
36. *Select > ‘Nuclei’ as smaller identified objects (Output from Identify Primary Object).*
37. *Select > ‘Yes’ to shrink smaller object prior to subtraction.* **Note:** Nuclei will now be subtracted from Full Doughnut area to produce a ring shaped ‘True Doughnut’ in the peri-nuclear region of the identified SN4741 cells. By selecting ‘Yes’ to shrink the smaller object the nucleus is shrunk by 1 pixel before subtracting the objects. This ensures a tertiary object is produced. Measurements such as area and fluorescence intensity of the segmented area can be acquired subsequently.

### Step 5: Identify Secondary Objects: SN4741 Cell Body

Following the identification of SN4741 nuclei using ***Identify Primary Object*** module the ***Identify Secondary Object*** module is used again. However, this time the module and method must find the edge of SN4741 cell bodies as secondary objects using cell body-specific Cy3 staining intensity. Again, this module involves a thresholding step which assigns pixels in the input image to foreground and background. To identify cell body:

38. *Select > ‘****Identify Secondary Objects’*** *module.*
39. *Input Image > Selects ‘CorrCy3’.*
40. *Select > Input Objects ‘nuclei’ (identified in* ***Identify Primary Object*** *module).*
41. *Input > ‘CellBody’ as name for identified objects.*
42. *Select > ‘Propagation’ as method to identify secondary objects.*
43. *Select > ‘Automatic’ as threshold method.*
44. *Input > ‘0.00’ as regularization factor.*
45. *Select > ‘No’ to fill holes in identified objects.*
46. *Select > ‘Yes’ to discard objects touching the image border.*
47. *Select > ‘Yes’ to discard associated primary objects.*
48. *Select > ‘No’ to retain the outlines of the identified secondary objects.*

### Step 6: Identify Tertiary Objects: SN4741 Cytoplasm

Similarly, to that described in ‘Step 4’ of MMP Protocol, the ***Identify Tertiary Objects*** module subtracts the shape of the smaller object (nucleus) from that of the larger identified object (i.e. cell body). Once removed the segmented area identified is the cell cytoplasm.

To identify Mitohealth stained SN4741 cytoplasm using Cell Profiler modules:

49. *Select >* ***Identify Tertiary Objects*** *module.*
50. *Select > ‘Cell Body’ as input for larger identified objects.*
51. *Select > ‘Nuclei’ as input for smaller identified objects.*
52. *Input > ‘Cytoplasm’ as name for tertiary objects to be identified.*
53. *Select > ‘Yes’ to shrink smaller object. Ensuring a cytoplasm is created for each nucleus.*
54. *Select > ‘No’ to Retain outlines of tertiary objects as these will not be utilised downstream.*

### Step 7: Enhance Mitochondrial Speckles

The ***Enhance Or Suppress Feature*** module is a CellProfiler module that suppresses or enhances selected image features such as speckles, ring shapes and neurites to improve the subsequent object identification by an ‘Identify module’. In this case it is used to enhance the punctate mitochondrial staining by applying image processing filters to the input image and giving a grayscale image output. Following the execution of this module *the ‘****Identify Primary Object’*** module is used again to identify mitochondrial puncta.

55. *Select > ‘CorrCy3’ as input image (image is illumination function corrected).*
56. *Input > ‘FilteredRed’ (or other user-defined name) for output grayscale image.*
57. *Select > ‘Enhance’ as operation type.*
58. *Select > ‘Speckles’ as feature type to be ‘Enhanced’.*
59. *Input > ‘20’ as feature size.*

### Step 8: Identify Primary Objects: Mitochondrial Fragments

This step utilises the **Identify Primary Objects** module to identify the mitochondrial puncta in the images enhanced in step 5. To identify mitochondrial puncta in enhanced x40 objective Cy3 channel images of MitoHealth stained SN4741 cells:

60. *Select >* ***Identify Primary Objects module.***
61. *Select > FilteredImage (output image from previous module) as input image.*
62. *Input > ‘Puncta’ (or other user-defined name) to assign to identified objects.*
63. *Select object diameter range > Min= 2; Max= 35.*
64. *Select > ‘Yes’ to discard objects outside the diameter range.*
65. *Select > ‘Yes’ to discard objects touching the bored of the image.*
66. *Select > ‘Per Object’ as Thresholding strategy and ‘Otsu’ as thresholding method.*
67. *Select > ‘Cytoplasm’ (previously identified tertiary objects) as masking objects.*
68. *Select > ‘Three classes’ thresholding and minimize the ‘weight variance’.*
69. *Assign> Middle intensity pixels to ‘Foreground’.*
70. *Select > ‘Automatic’ smoothing method for thresholding.*
71. *Input > ‘1.0’ as Threshold correction factor with ‘0.001’ lower and ‘0.005’ as upper bounds on threshold.*
72.
73. *Select > ‘No’ to automatically calculate size of smoothing filter for object de-clumping.*
74. *Input > ‘5’ as size of smoothing filter.*
75. *Select > ‘No’ to automatically calculate allowed distance between local intensity maxima.*
76. *Input > ‘5’ to suppress local maxima that are closer than this minimum allowed distance.*
77. *Select > ‘No’ to retain outlines of identified puncta.*
78. *Select > ‘Never’ to fill holes in identified objects.*
79. *Select > ‘Continue’ to handling of excessive number of identified objects-as puncta are small and therefore the number of puncta identified per image is usually high.* **Note:** At this stage in construction of CellProfiler analysis pipeline we have identified nuclei, cell body, mitochondrial fragments, cytoplasmic areas and peri-nuclear regions. Now that these cellular compartments have been identified using this CellProfiler pipeline, we must select which measurements and data to extract from our digital images using a set of measurement modules.

### Step 9: Measure Object Size and Shape

Although the ***Measure Object Size and Shape*** module can measure a vast array of shape and area features, we use this module to measure object size (area in pixels) of each cells’ mitochondrial puncta, cytoplasm and True Doughnut respective to CellProfiler pipeline.

To measure size and shape of object using ***Measure Object Size and Shape*:**

80. *Select >* ***Measure Object Size and Shape*** *module.*
81. *Select > ‘Cytoplasm’ or ‘Puncta’ or ‘True Doughnut’ or any combination of identified objects.* **Note:** Specific shape or size measurements (e.g. area) to output into spreadsheet is defined in a later module. This module simply selects which objects the measurements should be taken from.

### Step 10: Measure Object Intensity: True Doughnut or Cytoplasm

To measure the intensity of identified objects we use the ***Measure Object Intensity*** Cell Profiler module. To do so;

82. *Select > ‘CorrCy3’ illumination corrected image as image to measure.*
83. *Select > ‘True Doughnut’ or ‘Cytoplasm’ (or other object) identified in previous modules to measure.* **Note:** CorrCy3 image, as opposed to a Raw Image, is selected to acquire intensity measurements as this image previously undergoes illumination correction during the pre-processing stage. Specific object intensity measurements (e.g. mean, median, minimum, maximum intensity) to output into spreadsheet is defined in a later module. This module simply selects which objects the measurements should be taken from.

### Step 11: Relate Objects: Assign Mitochondrial Fragments to Cell Cytoplasm

The ***Relate Objects*** CellProfiler module is used here to assign a relationship between ‘child’ objects (e.g. puncta) with ‘parent’ objects (e.g. cytoplasm). This module can be utilised in order to count the number of puncta per SN4741 cell cytoplasm and calculate a mean measurement value for all ‘children’ objects which are associated with each ‘parent’ object. Objects (e.g. mitochondrial puncta) are considered ‘children’ of a ‘parent’ object (e.g. a cell cytoplasm) if the ‘child’ object is found within or at the edge of a ‘parent’ object. To assign the relationship between mitochondrial puncta and a cell cytoplasm:

84. *Select > ‘Puncta’ as input child objects (identified in second* ***Identify Primary Objects*** *module).*
85. *Select > ‘Cytoplasm’ as input parent objects (Identified in Identify Tertiary Objects module).*

### Step 12: Mask Objects to Telate Puncta to Cytoplasm of Origin

The ***Mask Objects*** module hides regions of the input image which are not regions of interest. This aids in the assignment of puncta to a cell cytoplasm of interest and allows for cell-by-cell puncta counts. In this example only SN4741 cell cytoplasm is selected as a region of interest and subsequently aids in exporting the Parent-Child (Cytoplasm-Puncta) data in the ***Export to Spreadsheet*** module which follows. To create a mask from previously identified objects (i.e cytoplasm):

86. *Select > ‘Filtered Red’ (the output of* ***Enhance or Suppress Feature*** *module) as input image.*
87. *Input > ‘Mask Red’ as (user defined) name for output image.*
88. *Select > ‘objects’ and ‘Cytoplasm’ to select object (cytoplasm) for mask.*
89. *Select > ‘No’ to invert mask to ensure mask is produced from the foreground area within the masking objects (cytoplasm).*

### Step 13: Export to Spreadsheet: Output Data into MS Excel Spreadsheet

The final module in these CellProfiler protocols is the Export to Spreadsheet. Here we assign a folder/file destination and name for the output data.

90. *Select > ‘Elsewhere’ and then ‘browse’ for desired output file location.*
91. *Select > ‘Yes’ to add prefix to file output file name.*
92. *Input > ‘Name’ (user-defined) prefix for output filename (e.g. MitoMMP or MitoPuncta).*
93. *Select > ‘No’ to overwrite without warning (Does not affect protocol-decided by end user).*
94. *Select > ‘No’ to add metadata columns to object data file (Does not affect protocol).*
95. *Select > ‘Yes’ to limit output to a size that allowed in Excel. If ‘no’ is selected, then upon exceeding limit on Excel spread sheet the pipeline will be terminated without data outputted.*
96. *Select > ‘NaN’ to represent data points which are numerical values are ‘infinite’ or ‘undefined’ in output spreadsheet.*
97. *Select > ‘Yes’ to select the measurements to export. The user can define which data are outputted to spread sheet. By selecting ‘No’ all data types will be outputted.*
98. Select the following data outputs from drop down box ‘select measurement to export’.

*To export number of nuclei (cell count) and puncta per image:*

99. *Image > Count > Nuclei;*
100. *Image > Count > puncta.*

*To export area of the objects True Doughnut and Cytoplasm:*

101. *True Doughnut > Area Shape > Area;*
102. *Cytoplasm > Area Shape > Area.*

*To export intensity of Mitohealth staining intensity in SN4741 peri-nuclear area:*

103. *True Doughnut > Intensity > Mean Intensity.*

*To export cell-by-cell puncta counts:*

104. *Cytoplasm > Children > Puncta > Count.*
105. *Select > ‘Yes’ to calculate the per-image mean values for object measurement. CellProfiler will calculate mean intensity, mean area, mean number of puncta per cell etc. for each image will then be exported into spreadsheet. Median and standard deviation measurements per image can also be calculated if desired.*
106. *File > Save Project As > input user defined project name. to save the CellProfiler image analysis pipeline for future use.* **Note:** CellProfiler protocol can now be opened and used on image set. To allow modules to run in sequence in an automated fashion select:
107. *File > Analyze Images.* A spreadsheet will be populated and saved as an output upon termination of all executable algorithms/modules in the Cell Profiler pipeline. This pipeline can be applied to experimental repeats.

Once these CellProfiler pipelines (CellProfiler Pipeline: Measuring Mitochondrial Membrane Potential and CellProfiler Pipeline: Measure SN4741 Mitochondrial Fragmentation) have been constructed it is then possible to automate image analysis for hundreds or thousands of images.

Below is a table (Table 2) containing the names and functions of the CellProfiler algorithm modules used in the above protocol(s).

**Table 2:** Cell Profiler Modules and Functions.

| **Cell Profiler Module** | **Function** |
| --- | --- |
| Correct Illumination - Calculate | Calculates an illumination function that is used to correct uneven illumination/lighting/shading or to reduce uneven background in images. |
| Correct Illumination - Apply | Applies an illumination function, usually created by Correct Illumination Calculate, to an image to correct for uneven illumination (uneven shading). |
| Identify Primary Objects | Identifies biological components of interest in grayscale images containing bright objects on a dark background (e.g. nuclei or speckles). |
| Identify Secondary Objects | Identifies objects (e.g., cell body) using objects identified by another module (e.g., nuclei) as a starting point, or seed. |
| Identify Tertiary Objects | Identifies tertiary objects (e.g., cytoplasm) by removing smaller primary objects (e.g. nuclei) from larger secondary objects (e.g., cell body), leaving a ring shape. |
| Enhance Or Suppress Features | Enhances or suppresses certain image features (e.g. speckles, ring shapes, and neurites), which can improve subsequent identification of objects (such as mitochondrial fragments). |
| Measure Object Size Shape | Measures several area and shape features of identified objects. |
| Measure Object Intensity | Measures several intensity features for identified objects. |
| Relate Objects | Assigns relationships; all objects (e.g. speckles) within a parent object (e.g. nucleus) become its children. |
| Export To Spreadsheet | Exports measurements into one or more files that can be opened in Excel or other spreadsheet programs. |

## Commentary

### Basic Protocol 1

#### Selection of an Appropriate Exposure Time

Prior to image acquisition ensure that time is taken to select the optimal exposure time as this will avoid saturation and a lack of dynamic range in the image acquired. To minimise potential of image saturation: i) maintain image acquisition conditions such as exposure times, lamp, and filters used etc. throughout the image acquisition process; ii) allow sufficient time for light source (lamp) to warm-up before running samples to avoid overexposure as lamp warms; an LED provides more consistency so use this as light source if possible; iii) Aim to quantitatively compare signals across the set of images to ensure that image pixel intensities do not become saturated due to increased fluorescence signal that may result from particular treatment regimens - for example positive experimental controls. To mitigate against this, aim to set the exposure time such that the resulting images use as much of the dynamic range of the camera as possible, but without saturating any images. For example, setting the image maximum to be ∼50-75% of the dynamic range is a good precaution and will allow for some images in the set being brighter than average without becoming saturated (Brown, 2007).

### Basic Protocol 2

#### Upload Image set, Image Sorting and Image Pre-processing and Illumination Correction

A limitation of many High-Content Screening (HCS) platforms, such as the InCell Analyser 2000, is that a flat-field correction (FFC) function cannot easily be implemented into the imaging process. There is the option to modify the images taken during a post-processing stage; however, this leaves the quality control of the FFC to the imaging software without any further input or quality control from the end user. Without a FFC application we can see that the intensity of an image varies widely from the centre to its edges (illustrated in figure 3). This spread of illumination intensity within an image will deter the accuracy of downstream image analysis. For example, regions of a cell body may appear shaded and therefore segmentation of image into biologically-relevant compartments (i.e. identification of cell body boundary) may be prevented.

The illumination pattern will change if different staining reagent(s) is used for a batch of samples or if there is a change in microscope components or setting of the optical path. For example, the illumination pattern may change throughout the same day or analysis as the lamp changes temperature. Therefore, in this protocol illumination is measured and correction applied for each image channel - that is Hoechst and Cy3. This illumination pattern must be corrected to ensure correct and comparable measurements in intra- and inter-image analysis. CellProfiler software has built-in modules to deal with these issues. The ***CorrectIlluminationCalculate*** module is used to measure the intensity pattern of a single or set of images and produces an intensity function which is to be applied across the entire image set. The ***CorrectIlluminationApply*** module then applies the calculated intensity function to the images prior to analysis. Correction of intensities can be important for segmentation and intensity measurement functions. In this method of illumination correction, we create and save an illumination function and apply this correction to images before segmentation. This can be performed using a separate Cell Profiler pipeline to create an ‘average’ illumination function for a specific fluorescence channel in a data set. Alternatively, as it is performed in this instance, the pre-processing (illumination correction) can be implemented at the start of an image analysis pipeline. This illumination function is saved as an image and is applied to the entire data set. The drawback of implementing illumination correction as part of a working pipeline, especially when dealing with large quantities of image files, is that calculation of the illumination function can add a considerable amount of time to the automated analysis process (Lindblad & Bengtsson, 2001).

#### Identification of Nuclei

*See ‘In-vitro rotenone assay: cell density’ in troubleshooting section for considerations.*

#### Identifying Secondary Objects: SN4741 Cell Body

Abnormal fragmentation of mitochondria is a trait observed in cells derived from patients afflicted with neurodegenerative diseases such as PD (Pieczenik & Neustadt, 2007). Pesticides such as rotenone are commonly used to generate PD models. In in-vivo rat models, systemic exposure to rotenone induces neurochemical, behavioural and neuropathological features of PD (Betarbet et al., 2000). These PD-like features are inclusive of nigrostriatal dopaminergic degeneration which is associated with the behavioural traits of hypokinesia and rigidity. The systemic exposure of rotenone in this rat model also resulted in other PD-associated neurochemical hallmarks such as the accumulation of ubiquitin and alpha-synuclein containing cytoplasmic inclusions (Betarbet et al., 2000). Moreover, evidence shows that cells exposed to rotenone, a mitochondrial Electron Transport Chain (ETC) complex-I inhibitor, results in a fragmented mitochondrial morphology *in-vitro*. An affect which is appears to be dependent on metabolic energy sensor, AMPK (Toyama et al., 2016).

Here we describe a protocol which allows for identification of entire mitochondria in SN4741 cell body and subsequent quantification of mitochondrial fragments and measurement of fragment size. This CellProfiler pipeline utilises aspects of the CellProfiler ‘Speckle Counting’ example pipeline with alterations to complement the analysis of SN4741 cells for mitochondrial fragments. The basic pipeline is similar to that described above, however, there are adjustments at i) ‘step 3’ as CellProfiler software is tweaked to identify SN4741 cell body; and ii) ‘step 4’ as the edges of entire SN4741 cell cytoplasm are identified as opposed to only the ring shaped ‘True Doughnut’ in the peri-nuclear region of the cell. Alterations are described in a step-by-step method below. During step 5 Note: This module uses ‘propagation’ as the method to identify dividing lines between clumped cell bodies (secondary objects) where there is a local change in staining intensity (i.e. cell bodies having different staining intensities; and foreground and background have different staining intensities). This algorithm varies from the watershed method (used in Image J macro) as the dividing lines between objects (cell bodies) are determined by both the changes in local image intensity (intensity gradient) and distance to the nearest primary object (nucleus). Dividing lines are placed where the image local intensity changes perpendicular to the boundary (Jones, Carpenter, & Golland, 2005). Therefore, this method is considered an improved method of object segmentation. Since ‘propagation’ is selected a user-defined regularization factor (given the symbol λ) must be input to inform the execution of the propagation algorithm. Propagation takes 1) the distance to nearest primary object and 2) intensity of the secondary object image into account when defining the dividing lines between objects. The regularization factor is used to decide the balance and weight placed on each factor during propagation and can be anywhere in the range from 0 to infinity. If a regularization value >1 is selected, the image intensity is almost completely ignored. Whilst selecting a regularization value of 0, means that the distance to the nearest primary object is ignored and propagation relies on image intensity. The larger the regularization value the more image intensity is ignored and dividing lines drawn become more reliant on distance to the nearest primary object.

#### Enhancing Mitochondrial Speckles

During this step the enhance operation is used to create an output image which predominantly contains user defined features. In this pipeline, the module produces a grayscale image largely composed of mitochondrial puncta. Feature size was determined by measurement of mitochondrial puncta using the ***Tools > Measure length*** CellProfiler function. CellProfiler suggests that when enhancing (or subtracting) the largest feature size in the image should be selected. In these MitoHealth images the largest mitochondrial puncta measured were approximately 20 pixels in diameter. Speckle feature type was selected to enhance mitochondrial puncta in each image. CellProfiler defines a speckle as an area of enhanced pixel intensity relative to its immediate neighbourhood using a ‘tophat’ filter. This filter used grayscale image erosion within a set radius (Approx. object diameter = 20 pixels; therefore radius = 10 pixels), and then dilation. The speckles are enhanced by this module making them easier to identify with the ***Identify Primary Object*** module. See figure 9 (e-g) for example of identified puncta.

**Figure 9:**
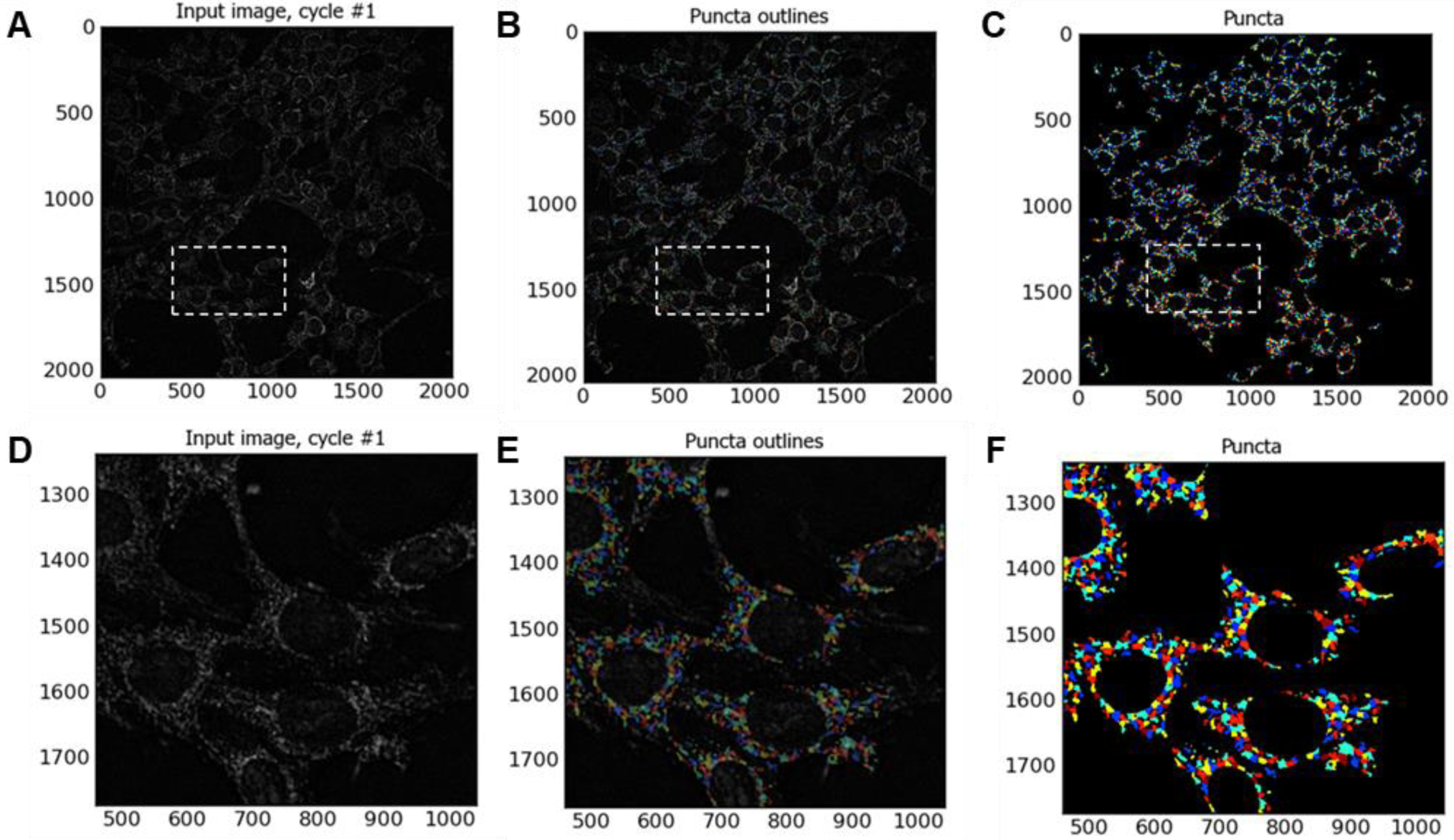
Identification of Mitochondrial Puncta. Images illustrating mitochondrial puncta identified with automated image analysis CellProfiler pipeline. Image (**A**) is an example input image to the Identify Primary Objects module. (**B**) illustrates the outlines of identified mitochondrial puncta, and (c) shows object masks of identified mitochondrial puncta. Dashed boxes on (**A**), (**B**) and (**C**) indicate where CellProfiler zoom function has been utilised to generate images (**D**), (**E**) and (**F**), respectively, for illustrative purposes.

#### Identifying Mitochondrial Fragments as Primary Objects

In identifying mitochondrial fragments in step 8, we use ‘Per object’ thresholding strategy which relies on the identification of a cellular compartment in a previous CellProfiler module. In this example, SN4741 cell ‘Cytoplasm’ is identified by ***Identify Tertiary Object*** module. Using this strategy means each individual cell cytoplasm has its own applied threshold. Pixels outside of the object are considered background. CellProfiler suggests this as a useful method for identifying particles in cellular sub-compartments if staining varies between objects/cell-to-cell. Otsu method of thresholding (named after Nobuyuki Otsu) calculates image threshold by minimising the variance within the pixel that are considered to be classed and assigned to foreground or background (Otsu, 1979; Sankur, 2004). CellProfilers’ application of Otsu thresholding method can be applied to assign pixels to either two-classes (foreground and background), or three-classes (foreground, mid-ground and background). Two-class is selected as foreground (regions of interest i.e. puncta) background are easily distinguishable following enhancement module.

## Critical Parameters and Troubleshooting

The in-vitro rotenone assay described here is a 2D model with 2D image acquisition. We recognise that that 3D culture of neuronal spheroids and organoids may be a more physiologically relevant to studying the neurodegenerative disease and modelling cellular interactions within the human brain (Chlebanowska, Tejchman, Sułkowski, Skrzypek, & Majka, 2020). In our laboratory we sought to develop an easily reproducible and resource efficient method for the screening of multiple compounds for their potential neuroprotective effects in models of PD using readily available equipment, skills and resources. We also recognise the use of live-cell-imaging techniques and approaches to answer key questions regarding the mechanisms or potential treatments of neurodegenerative diseases (Bakota & Brandt, 2009). In our protocol described in this paper we utilise paraformaldehyde fixation to immortalise the phenotype/morphology under investigation– to this end we use fluorescence probes which are compatible with PFA fixation process. Fixation of our sample also allows for the option of multiplexing with antibody immunocytochemistry. We have a simple easily replicable step-by-step guide to executing the protocol.

Basic Protocol 1 and Basic Protocol 2 provide comprehensive step-by-step guidance to for execution of this protocol. Additional considerations and troubleshooting notes are described below.

### In-Vitro Rotenone Assay: Cell Density

The protocol in basic protocol 1 describes the seeding of 3,000 cells in each treatment well. SN4741 cells under described culture conditions have a cell doubling time of approximately 23hrs. The timeline from cell seeding through to experimental endpoint is 72hr. Seeding cells at 3,000 cells per well results in cell density at experimental endpoint of approximately 80%. Thus, mitigating against over confluent cell density. Over-confluent cells at experimental endpoint means that downstream image analysis difficult and can result in inaccuracies in image segmentation– specifically nuclear segmentation, where the goal is to identify discrete nuclei to build and execute subsequent image analysis steps. Misidentification of nuclei may result in repercussions for the subsequent image analysis steps and inaccuracies in data measurement. See step 2 in basic protocol 2 for nuclei segmentation.

### In-Vitro Rotenone Assay: Rotenone Efficacy

On occasions we have observed a loss of efficacy of rotenone. Resulting in challenge resulting in non-significant loss of SN4741 neurons following 24hr. In order to mitigate against this, the reconstituted rotenone (in DMSO) is stored at −4°C protected from light and air using parafilm or tape to seal aliquots. We recommend aliquoting small volumes (10uL) of rotenone for storage and a max of 2-3 freeze-thaw cycles. There is also a possibility that rotenone falls out of aqueous solution with DMSO on freeze-thaw cycle. In light of this, we highly recommend through mixing of rotenone aliquots by vortex prior to use.

### Sample Storage Considerations

Due to unforeseen circumstances it might not be possible to acquire images for samples immediately following staining procedure as recommended by manufacturers. Should this problem arise, place samples in cold room (4°C) overnight and protect from light; should the sample need to be kept longer Prolong Gold™ (Invitrogen, cat no. P36930) should be used following manufacturer’s instructions to retain sample fluorescence integrity, protect from light and store at 4°C. Ensure that the sample plate is left to acclimatise to room temperature prior to imaging to avoid condensation.

### Data Storage Considerations

Images (.tiff, 16-bit) acquired using the automated fluorescence microscopy system are saved to an external hard drive. Each .tiff image acquired in protocol requires approximately 8MB of storage. The main consideration here is ensuring that storage device has sufficient storage capacity for experimental setup. For more information on digital image data see ‘Digital Image Data’ in Supplementary Protocol Information.

### Time Considerations

Outline of basic protocol 1 and basic protocol 2 experimental procedure and associated timelines can be seen in Figure 10. Figure 11 illustrated the experimental timelines of the in-vitro rotenone PD model.

**Figure 10.**
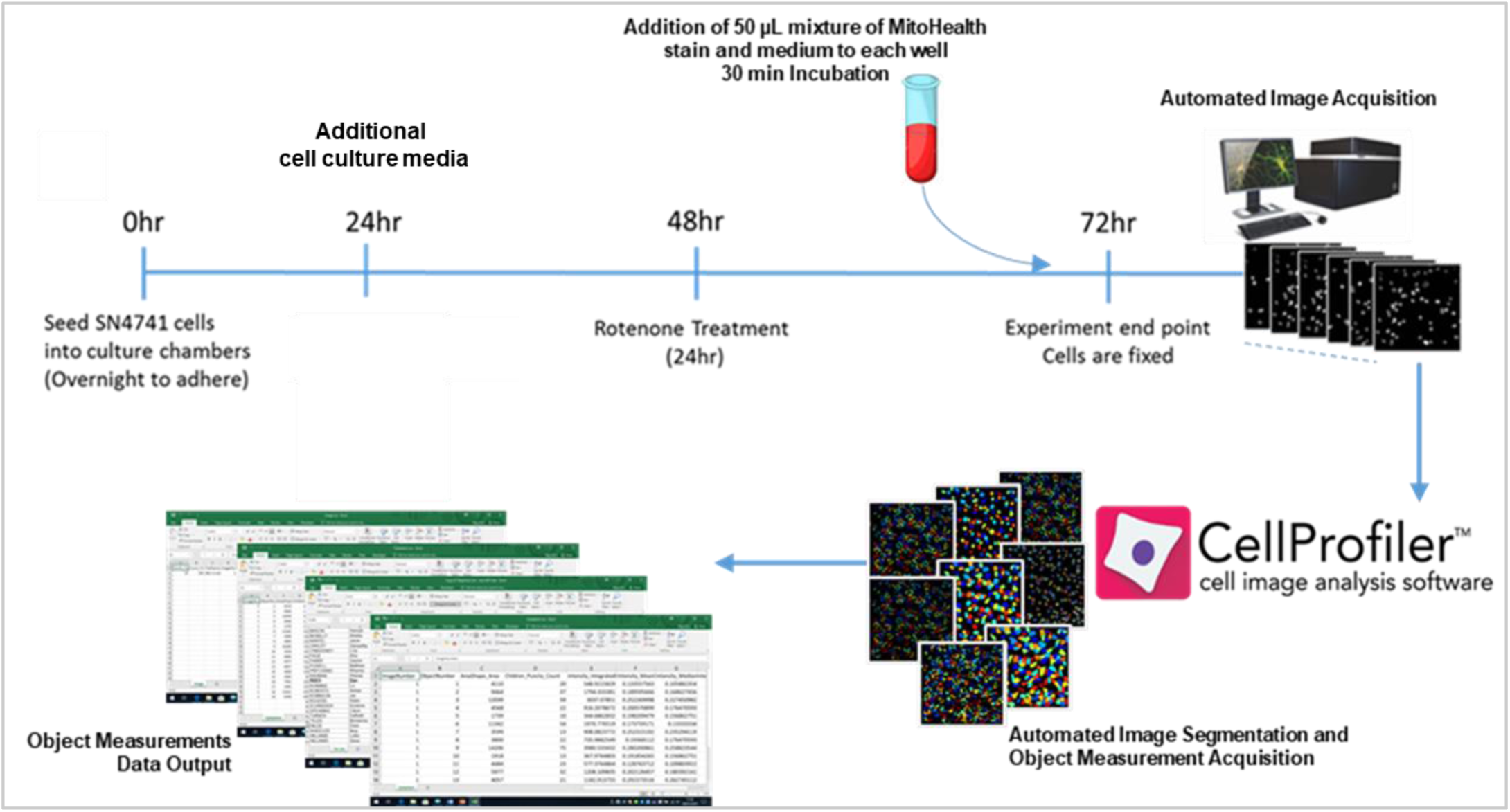
Experimental Outline and Associated Timelines. This diagram illustrates the outlines key steps of basic protocol 1 and basic protocol 2. This includes wet-laboratory preparation of SN4741 cells; cell staining process; automated image acquisition; and execution of CellProfiler image analysis software for image segmentation and data output.

**Figure 11.**
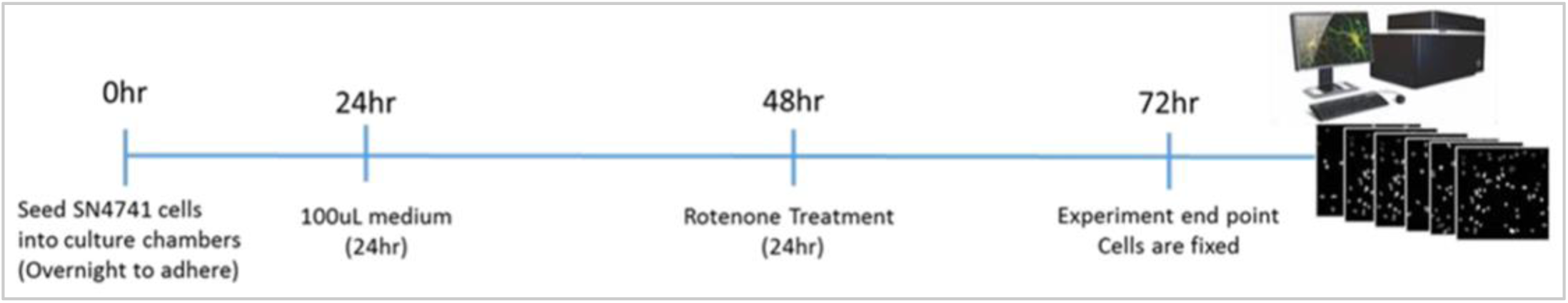
Basic Protocol 1 Timeline. Illustration of Basic Protocol 1 timeline including cell seeding; addition of cell culture medium following 24hr incubation; 24hr rotenone challenge, and wet laboratory experimental endpoint. Cell staining is executed at 71.5hr into experimental timelines - not illustrated in this diagram.

**Figure 12.**
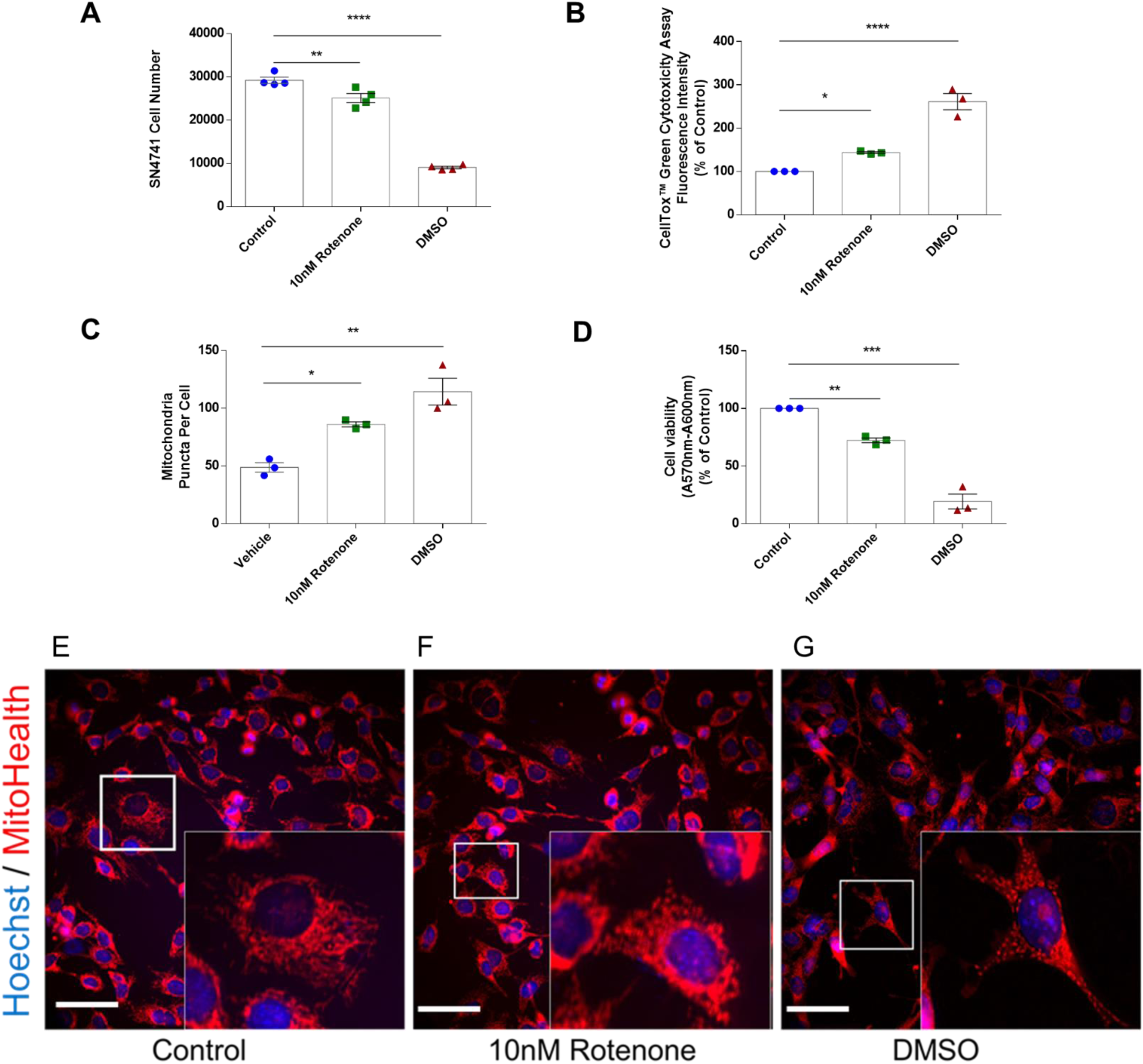
Rotenone-Based Parkinson’s disease model results in Cell Loss, Decreases Viability and Increases SN4741 Cell Cytotoxicity and Mitochondrial Fragmentation. Figure 1 (**A**), (**B**), (**C**) and (**D**) represents SN4741 cell count, cytotoxicity, mitochondrial fragments per cell, and viability data respectively. A decrease in cell number (**A**); increase in cell cytotoxicity (**B**) and mitochondria fragments per cell (**C**); and a decreased cell viability (**D**) is observed following 24hr challenge with 10nM rotenone. SN4741 cell number (A) is expressed as mean cell count from cells treated with in quadruplicate (n=4); SN4741 cytotoxicity, mitochondrial fragmentation and viability data (**B** - **D**) expressed as mean cell count from cells treated in triplicate for 3 experimental repeats (n=3). Cell Cytotoxicity (**B**) and viability data (**D**) are expressed as percentage of control (0nM). Statistical analysis was performed by One-way ANOVA with Dunnett’s post-hoc analysis (**A** - **D**) in graph pad prism 6 software, with P<0.05 (*), P<0.01 (**), P<0.001 (***) considered significant when compared to treatment control (0nM Rotenone). Diagrams **E, F** and **G** are representative merged pseudo-coloured DAPI and Cy3 images for Control, 10nM rotenone and DMSO challenged SN4741 cells, respectively. Images (**E**-**G**) represent x40 magnification images with scale bar = 40µm. Each merged image has a representative enhance image for graphical purposes highlighted by a white box. The enhanced regions shown in each merged image (**E**-**G**) defines a region of the image which have been enhanced for illustration of fragmentation morphology under treatment conditions.

Another consideration is estimated run time on each image set (1 DAPI image and 1 Cy3 Image). In this CellProfiler pipeline is typically 1-1.5 minutes. Run time for 10 image sets (10 DAPI and 10 Cy3 images) is approximately 14 minutes with step-by-step display in CellProfiler software enabled; and 9.5 mins with display disabled. Run time for 52 images is approximately 1hr 5mins with display and 48 mins without display. Display can be toggled on/off for each module in CellProfiler software pipeline panel. Run times represent those achieved using a Toshiba TECRA A50-C-218 with Inter(R) CoreTM i7-6500U CPU @2.5GHz to run this image analysis protocol. Run time will be quicker on laptops or desktop computers with additional RAM and CPU.

### Software for Statistical Analysis

The terminating step in the CellProfiler pipeline is exporting measurements and data. In this protocol data is exported as comma separated value (.csv) files which can be opened in MS Excel. Graphing and biostatistics can be performed in MS Excel or commercially available software such as Graphpad Prism or MATLAB. Statistical analysis of mean puncta number per cell, mean fluorescence intensity per cell and cell number has been performed by One-way ANOVA with Dunnett’s post-hoc analysis, with P<0.05 (*), P<0.01 (**), P<0.001 (***) considered significant when compared to control (0nM Rotenone).

### PPE and COSHH Considerations

Rotenone and PFA pose a hazard to human health and will need to be disposed of appropriately. Relevant good laboratory practise, PPE and COSHH guidelines must be adhered when handling these chemical and compounds. In light of this appropriate risk assessments should be performed and approved by the organisation’s Health & Safety committee.

### Additional Applications for Image Analysis Pipeline

The simplicity of this protocol means that it is easily adapted to the identification and quantification of other sub-cellular organelles. An example of such applications includes identification of fluorescently lebelled lysosomes and autophagosomes from 2D images acquired through HCI in the analysis of autophagy. The laboratory has found use for this pipeline in a myriad of cell types including HT22, SN4741 and differentiated and undifferentiated human ReN cells with slight modification of parameters such as ‘typical nucleus diameter’ in ***Identify primary objects*** module.

## Supplementary Protocol Information

### Digital Image Data

The protocols described in this paper perform data capture on digital images of biological samples in an uncompressed 16-bit TIFF format. Details on digital image file displayed in table below:

| IN Cell Analyzer 2000 Output - Digital Image Data |  |
| --- | --- |
| • File size= 7.93 MB (8,323,678 bytes = 66,585,600 bits*)<br>• Width= 2040 pixels<br>• Length= 2040 pixels<br>• Horizontal Resolution= 68649 dpi<br>• Vertical Resolution= 68649 dpi | • Bit-Depth= 16-bit<br>• Compression= Uncompressed<br>*\* Length \* Width \* Bit-Depth = Image Data Size (bits)* |

Images captured on the IN Cell Analyzer 2000 (or equivalent HCS system) should be saved in an uncompressed (lossless) format. Here we capture 16-bit 2040 x 2040 pixel images on an external hard drive for image analysis purposes.

**Note:** Lossless compression algorithms reduce file size while preserving a perfect copy of the original uncompressed image containing the detail and full pixel range/bit-depth available when stored as an image of 16 bit-depth. The levels (depth) of colour (referred to as grey scale in binary images) is important for effect collection of numerical values (a specific example is intensities) in the image analysis process. Lossless compression (or uncompressed files) usually result in larger files, whereas lossy compression of files generally produce smaller files but at the expense of image quality. Files which are saved can be converted to compressed files at the expense of image quality; however, it is not possible to retrieve lossless image quality (to ‘original’) from an image which has been saved in a compressed method. It is recommended that uncompressed 16-bit TIFF format files are utilised for protocols which refer to a HCS system, such as the IN Cell Analyzer, as the method of image acquisition.

**Note**: Bit depth describes the number of data bits available to represent the intensity value of a single pixel. It is also known as bits per pixel (bpp). An image file format’s bit depth indicates the number of separate grayscale intensity values (graylevels) that are allowable by the file format:

→ 8-bit images have 2^8 available pixel intensities, with a range of 0-255
→ 12-bit images have 2^12 available pixel intensities, with a range of 0-4095
→ 16-bit images have 2^16 available pixel intensities, with a range of 0-65535

CellProfiler normalizes (contrast-stretches) images from raw image files. On most imaging platforms 12-bit images are saved as a 16-bit images and it is common when visualising raw images on standard computer software (e.g. MS media player) that your images will appear a dark when presented raw. CellProfiler analysis uses the actual (raw) intensity values for image analysis and data acquisition and not the contrast stretched image which is observed in the CellProfiler for visualisation purposes.

### IN Cell Analyzer 2000 (GE Healthcare) Specifications

**Description:** The GE Healthcare IN Cell Analyzer 2000 is an automated cellular and subcellular imaging system for fixed and live cells. Like other automated high-content-screening (HCS) systems, the IN Cell 2000’s features such as CO_2_, humidity and temperature (heating) controls allow for the systems applicability to both fixed and live cell automated image acquisition. The systems allows both manual investigative microscopy as well as fully-automated HCS and the study of a variety of sample inputs, from organelles to cells, tissues, and whole organisms, and from fixed end-point assays to extended live-cell studies. The system is composed of a combination of proprietary optics, fast hardware and software autofocus, with a high-performance CCD camera. This results in rapid well-to-well imaging and light source exposure time in order to minimise compromise on image quality and cell health. The designed features and functions on this system and supporting software enables complex biological techniques to be incorporated with ease into your high content program. Applications performed with IN Cell Analyzer 2000 include compound screening, stem cell assays, phenotypic profiling, predictive toxicology, RNAi screening, tissue microarrays, whole organism imaging, signalling pathway analysis, neurite outgrowth/neuronal function, cell cycle studies, cell migration, micronucleus assay, co-localization analysis, automated slide imaging, organelle & protein trafficking, receptor activation, morphology analysis, DNA content analysis, apoptosis/cell viability, mitochondrial function, colony counting (e.g. stem cells) and fluorescent in situ hybridization.

#### Features of the IN Cell 2000 Analyzer (HCS System)

The IN Cell Analyzer 2000 system was used extensively to carry out image acquisition for the protocols described in this paper. The table below lists the IN Cell Analyzer 2000’s key features which allow for its application to a multiplicity of assays and analyses.

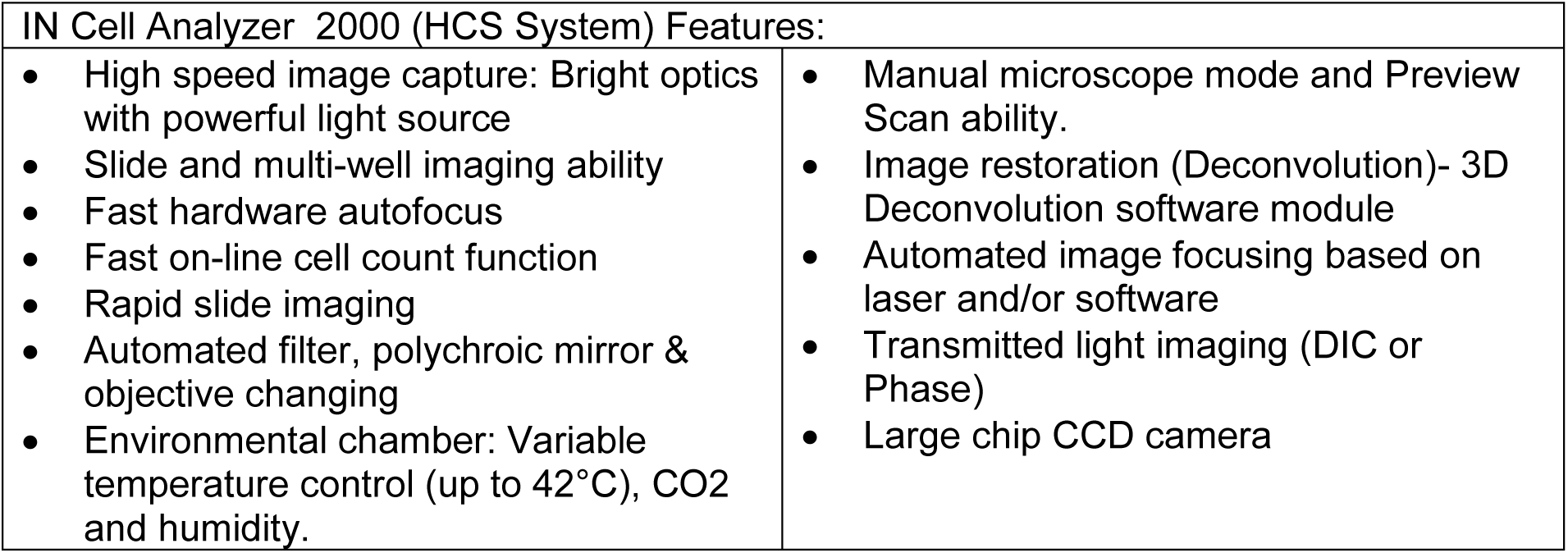

| IN Cell Analyzer 2000 (HCS System) Features: |  |
| --- | --- |
| • High speed image capture: Bright optics with powerful light source<br>• Slide and multi-well imaging ability<br>• Fast hardware autofocus<br>• Fast on-line cell count function<br>• Rapid slide imaging<br>• Automated filter, polychroic mirror & objective changing<br>• Environmental chamber: Variable temperature control (up to 42°C), CO<sub>2</sub> and humidity. | • Manual microscope mode and Preview Scan ability.<br>• Image restoration (Deconvolution)- 3D Deconvolution software module<br>• Automated image focusing based on laser and/or software<br>• Transmitted light imaging (DIC or Phase)<br>• Large chip CCD camera |

#### IN Cell 2000 Analyzer (HCS System) Microscope Objective Lenses, Mirrors and Filters

The IN Cell Analyzer is equipped with a range of objective lenses, mirrors and filters to facilitate its application to a multiplicity of experimental analysis scenarios and fluorescence probe detection. The details of objectives, mirrors & filters and related fluorescence excitation/emission detection ranges are listed in the tables below.

##### Objectives

| IN Cell Analyser 2000 Objectives |  |  |  |
| --- | --- | --- | --- |
| Magnitude | NA | ASAC (Range mm) | Working Distance |
| 2X | 0.1 | No | 8.5 mm |
| 10X | 0.45 | No | 4.0 mm |
| 20X | 0.45 | Yes (0-2) | 7.5 mm |
| 60X | 0.95 | Yes (0.11-0.23) | 0.15 mm |

##### Polychroic Mirrors & Filters

The IN Cell Analyzer 2000 imaging system comes complete with polychroic mirrors and filters covering all the most popular dyes and fluorescence proteins

| IN Cell Analyser 2000 Polychroic Mirrors & Filters |  |
| --- | --- |
| Polychroic | Suitable for |
| QUAD 1 | • DAPI/Hoechst/Quantum Dots<br>• eGFP/FITC<br>• Cy3/TRITC/DsRed/mOrange/Rhodamine/ dTomato<br>• Cy5/DRAQ5 |
| QUAD 2 | • DAPI/Hoechst/Quantum Dots<br>• eGFP/FITC<br>• Texas Red/HcRed1/mRaspberry/mPlum/mCherry<br>• Cy5/DRAQ5 |

## Notes

### Competing Interest Statement

The authors have declared no competing interest.

## Literature Cited

Bakota, L., & Brandt, R. (2009). Chapter 2 Live-Cell Imaging in the Study of Neurodegeneration. https://doi.org/10.1016/S1937-6448(09)76002-2

Betarbet, R., Sherer, T. B., MacKenzie, G., Garcia-Osuna, M., Panov, A. V., & Greenamyre, J. T. (2000). Chronic systemic pesticide exposure reproduces features of Parkinson’s disease. Nature Neuroscience, 3(12), 1301–1306. https://doi.org/10.1038/81834

Brown, C. M. (2007). Fluorescence microscopy - avoiding the pitfalls. Journal of Cell Science, 120(10), 1703–1705. https://doi.org/10.1242/jcs.03433

Chlebanowska, P., Tejchman, A., Sułkowski, M., Skrzypek, K., & Majka, M. (2020). Use of 3D Organoids as a Model to Study Idiopathic Form of Parkinson’s Disease. International Journal of Molecular Sciences, 21(3), 694. https://doi.org/10.3390/ijms21030694

Franco, R., Li, S., Rodriguez-Rocha, H., Burns, M., & Panayiotidis, M. I. (2010). Molecular mechanisms of pesticide-induced neurotoxicity: Relevance to Parkinson’s disease. Chemico-Biological Interactions, 188(2), 289–300. https://doi.org/10.1016/j.cbi.2010.06.003

Haque, M. E., Thomas, K. J., D’Souza, C., Callaghan, S., Kitada, T., Slack, R. S., … Park, D. S. (2008). Cytoplasmic Pink1 activity protects neurons from dopaminergic neurotoxin MPTP. Proceedings of the National Academy of Sciences, 105(5), 1716–1721. https://doi.org/10.1073/pnas.0705363105

Hoglinger, G. U., Lannuzel, A., Khondiker, M. E., Michel, P. P., Duyckaerts, C., Feger, J., … Hirsch, E. C. (2005). The mitochondrial complex I inhibitor rotenone triggers a cerebral tauopathy. Journal of Neurochemistry, 95(4), 930–939. https://doi.org/10.1111/j.1471-4159.2005.03493.x

Jayaraj, R. L., Tamilselvam, K., Manivasagam, T., & Elangovan, N. (2013). Neuroprotective Effect of CNB-001, a Novel Pyrazole Derivative of Curcumin on Biochemical and Apoptotic Markers Against Rotenone-Induced SK-N-SH Cellular Model of Parkinson’s Disease. Journal of Molecular Neuroscience, 51(3), 863–870. https://doi.org/10.1007/s12031-013-0075-8

Jones, T. R., Carpenter, A., & Golland, P. (2005). Voronoi-Based Segmentation of Cells on Image Manifolds. https://doi.org/10.1007/11569541_54

Kamentsky, L., Jones, T. R., Fraser, A., Bray, M.-A., Logan, D. J., Madden, K. L., … Carpenter, A. E. (2011). Improved structure, function and compatibility for CellProfiler: modular high-throughput image analysis software. Bioinformatics, 27(8), 1179–1180. https://doi.org/10.1093/bioinformatics/btr095

Lindblad, J., & Bengtsson, E. (2001). A comparison of methods for estimation of intensity nonuniformities in 2D and 3D microscope images of fluorescence stained cells. Proceedings of the 12th Scandinavian Conference on Image Analysis (SCIA), 264–271.

McQuin, C., Goodman, A., Chernyshev, V., Kamentsky, L., Cimini, B. A., Karhohs, K. W., … Carpenter, A. E. (2018). CellProfiler 3.0: Next-generation image processing for biology. PLOS Biology, 16(7), e2005970. https://doi.org/10.1371/journal.pbio.2005970

Mehta, S. L., & Li, P. A. (2009). Neuroprotective Role of Mitochondrial Uncoupling Protein 2 in Cerebral Stroke. Journal of Cerebral Blood Flow & Metabolism, 29(6), 1069–1078. https://doi.org/10.1038/jcbfm.2009.4

Otsu, N. (1979). A Threshold Selection Method From Gray-Level Histograms. IEEE Transactions on Systems, Management, and Cybernetics, 9(1), 62–66.

Padmanabhan, K., Eddy, W. F., & Crowley, J. C. (2010). A novel algorithm for optimal image thresholding of biological data. Journal of Neuroscience Methods, 193(2), 380–384. https://doi.org/10.1016/j.jneumeth.2010.08.031

Pieczenik, S. R., & Neustadt, J. (2007). Mitochondrial dysfunction and molecular pathways of disease. Experimental and Molecular Pathology, 83(1), 84–92. https://doi.org/10.1016/j.yexmp.2006.09.008

Pozo Devoto, V. M., & Falzone, T. L. (2017). Mitochondrial dynamics in Parkinson’s disease: a role for α-synuclein? Disease Models & Mechanisms, 10(9), 1075–1087. https://doi.org/10.1242/dmm.026294

Reddy, P. H. (2009). Role of Mitochondria in Neurodegenerative Diseases: Mitochondria as a Therapeutic Target in Alzheimer’s Disease. CNS Spectrums, 14(S7), 8–13. https://doi.org/10.1017/S1092852900024901

Sandhu, L. C., Warters, R. L., & Dethlefsen, L. A. (1985). Fluorescence studies of Hoechst 33342 with supercoiled and relaxed plasmid pBR322 DNA. Cytometry, 6(3), 191–194. https://doi.org/10.1002/cyto.990060304

Sankur, B. (2004). Survey over image thresholding techniques and quantitative performance evaluation. Journal of Electronic Imaging, 13(1), 146. https://doi.org/10.1117/1.1631315

Scorrano, L., Petronilli, V., Colonna, R., Di Lisa, F., & Bernardi, P. (1999). Chloromethyltetramethylrosamine (Mitotracker Orange TM) Induces the Mitochondrial Permeability Transition and Inhibits Respiratory Complex I. Journal of Biological Chemistry, 274(35), 24657–24663. https://doi.org/10.1074/jbc.274.35.24657

Scott, I., & Youle, R. J. (2010). Mitochondrial fission and fusion. Essays in Biochemistry, 47, 85–98. https://doi.org/10.1042/bse0470085

Sherer, T. B., Betarbet, R., Stout, A. K., Lund, S., Baptista, M., Panov, A. V., … Greenamyre, J. T. (2002). An In Vitro Model of Parkinson’s Disease: Linking Mitochondrial Impairment to Altered α-Synuclein Metabolism and Oxidative Damage. The Journal of Neuroscience, 22(16), 7006–7015. https://doi.org/10.1523/JNEUROSCI.22-16-07006.2002

Son, J. H., Chun, H. S., Joh, T. H., Cho, S., Conti, B., & Lee, J. W. (1999). Neuroprotection and Neuronal Differentiation Studies Using Substantia Nigra Dopaminergic Cells Derived from Transgenic Mouse Embryos. The Journal of Neuroscience, 19(1), 10–20. https://doi.org/10.1523/JNEUROSCI.19-01-00010.1999

Toyama, E. Q., Herzig, S., Courchet, J., Lewis, T. L., Loson, O. C., Hellberg, K., … Shaw, R. J. (2016). AMP-activated protein kinase mediates mitochondrial fission in response to energy stress. Science, 351(6270), 275–281. https://doi.org/10.1126/science.aab4138

